# Environmental exposures and familial background alter the induction of neuropathology and inflammation after SARS-CoV-2 infection

**DOI:** 10.1101/2024.12.02.626375

**Authors:** Debotri Chatterjee, Drishya Kurup, Richard Jay Smeyne

## Abstract

Basal ganglia disease has been reported as a post-infection sequela of several viruses, with documentation of this phenomenon from the H1N1 Spanish flu to the recent COVID-19 (SARS-CoV-2) pandemic. SARS-CoV-2 infection leads to multisystem deficits, including those affecting the nervous system. Here, we investigated whether a SARS-CoV-2 infection alone increases the susceptibility to develop parkinsonian phenotypes in C57BL/6J mice expressing the human ACE2 receptor, or in addition to two well-known toxin exposures, MPTP and paraquat. Additionally, we examined mice carrying a G2019S mutation in the LRRK2 gene. We also examined if vaccination with either an mRNA- or protein-based vaccine can alter any observed neuropathology. We find that the infection with the WA-1/2020 (alpha) or omicron B1.1.529 strains in ACE2 and G2019S LRRK2 mice both synergize with a subtoxic exposure to the mitochondrial toxin MPTP to induce neurodegeneration and neuroinflammation in the substantia nigra. This synergy appears toxin-dependent since we do not observe this following exposure to the direct redox-inducing compound paraquat. This synergistic neurodegeneration and neuroinflammation is rescued in WT mice that were vaccinated using either mRNA- and protein-based vaccines directed against the Spike protein of the SARS-CoV-2 virus. However, in the G2019S LRRK2 mutant mice, we find that only the protein-based vaccine but not the mRNA-based vaccine resulted in a rescue of the SARS-CoV-2 mediated neuropathology. Taken together, our results highlight the role of both environmental exposures and familial background on the development of parkinsonian pathology secondary to viral infection and the benefit of vaccines in reducing these risks.

## Introduction

The etiology of Parkinson’s disease is multifactorial. It is estimated that the vast majority of individuals manifest the disease due to a variety of environmental factors or other idiopathic mechanisms; while approximately 10-15% of PD cases are linked to a number of familial mutations^1–4^. The most common familial genes underlying PD include the leucine-rich repeat kinase 2 (LRRK2)^5,6^, glucocerebrosidase (GBA)^7^, and PTEN-induced kinase 1 (PINK1)^8^. Of the non-familial cases, the development of PD has been epidemiologically or experimentally linked to several environmental exposures, including toxins^9^ such as paraquat^10,11^ and rotenone^11,12^, as well as bacterial and viral infections^13^. One of the main hypotheses- at this time- is that these familial factors interact with environmental exposures to synergize effects, often called the “multiple-hit hypothesis” in PD^14–16^.

One environmental factor linked to the development of PD is viruses^17–22^. In fact, neurological sequelae of viral infections have been reported for much of recorded medical history^22,23^. However, the most famous of these cases was described by Constantino von Economo^24,25^, who reported a parkinsonian syndrome called encephalitic lethargica that was described as coincident with the 1918 Spanish influenza pandemic^26–28^. Since then, several other viral infections^29^, both neurotropic and non-neurotropic, have been associated with the development of basal ganglia disease. In particular, a number of strains of influenza, including H5N1 and H1N1 have been linked to an increased risk of PD in both pre-clinical models^16,30–32^ as well as large-scale case-control studies^29,33^. Proposed mechanisms underlying the induction of parkinsonism following influenza infection include both direct effects secondary to CNS infection^34^ as well as indirect action through signaling of peripherally-derived inflammatory cytokines ^20,35,36^.

In 2019, a novel beta coronavirus outbreak (Severe Acute Respiratory Syndrome Coronavirus 2, or SARS-CoV-2) was reported in Wuhan, China^37^. In the years since, it has infected over 700 million individuals, as well as caused over 7 million deaths in the pandemic known as COVID-19^38^. SARS-CoV-2 is a positive-sense, single-stranded RNA virus that finds entry through binding to Angiotensin-converting enzyme 2 (ACE2) receptors in the lung epithelia^39,40^. Similar to other coronavirus family members SARS-CoV-1 and MERS, it primarily causes respiratory disease with symptoms ranging from cough and difficulty breathing to pneumonia and ARDS (acute respiratory distress syndrome)^41^. Additionally, infection with this virus has been shown to trigger an inflammatory cytokine and chemokine response, further exacerbating its induced symptoms^42–45^. In addition to its respiratory effects, SARS-CoV-2 has also been known to affect several other organ systems, including the nervous system; with neurological symptoms including anosmia and ageusia, headache, stroke, seizure, cognitive difficulty and “brain fog”^46^. Additionally, SARS-CoV-2 infection has also been associated with development of both acute parkinsonism^47–51^, and exacerbation of motor and non-symptoms in diagnosed PD patients ^52,53^.

Since the first outbreak of SARS-CoV-2, this virus has rapidly mutated, with each different strain showing varying infectivity as well as different rates of mortality and morbidity. Peak mortality was seen following infection with the alpha (B.1.1.7) and delta (B.1617.2) variants, with lower mortality seen following infection with the omicron strain (B1.1.529) and its sub-variants^54,55^. Since the fall of 2020, commercially available vaccines against SARS-CoV-2 have proven very effective in reducing mortality from the virus. In the USA, the majority of vaccinations given so far have been one of two mRNA-based vaccines: the Moderna vaccine (brand name Spikevax)^56^ and the Pfizer-BioNTech vaccine (brand name: Comirnaty)^57^, which both showed about 95% efficacy in their phase 3 trials targeting the original B.1.1.7 alpha strain. The third vaccine in circulation is the Novavax vaccine (brand names Nuvaxovid and Covovax)^58^, a more traditional protein-based vaccine which has shown an efficacy of about 90% in its clinical trial.

In this study, we compared the neuropathological and immunological effects of the WA- 1/2020 (B1.1.1.7) and an omicron strain (B1.1.529) of SARS-CoV-2 as well as empirically determined if COVID-19 infection synergized with environmental agents that have been linked to parkinsonism (MPTP, paraquat) or a familial PD mutation (G2019S LRRK2) to exacerbate basal ganglia pathology. We also examined if vaccination against COVID-19 could reduce, or even eliminate, any increased pathology secondary to any observed synergistic effects. Here we report that both the WA-1/2020 and an omicron strain of COVID-19 equally induced basal ganglia pathology despite much lower mortality and morbidity associated with the omicron strain. We also show that mice heterozygous for the G2019S LRRK2 allele have significantly increased substantia nigra pars compacta (SNpc) dopaminergic (DA) neuron loss and neuroinflammation following either WA-1/2020 or omicron SARS-CoV-2 infection. This synergy appears to be agent specific since we found that MPTP^59^ but not paraquat, induces a SARS-CoV-2 infection synergistic effect^60^. We also show that this synergistic viral-induced neurodegeneration and inflammation is ameliorated by vaccination against SARS-CoV-2 infection using either mRNA-based or protein adjuvant-based technologies; however, the mRNA-based vaccine did not rescue viral-induced sensitivity in mice carrying a familial G2019S allele of LRRK2 mutation.

## Methods

### Animals

8-14 week old WT (C57BL/6J) (Jackson Laboratory, Bar Harbor, ME; cat#000664), B6.Cg-Tg(K18-ACE2)2Prlmn/J (ACE2, Jackson Laboratory, Bar Harbor, ME; cat# 034860) and C57BL/6-Lrrk2tm4.1Arte (G2019S LRRK2 ki/ki, Taconic, NY; cat# 13940) were used in this study. Approximately equal numbers of males and females were used in all experiments, and all controls were age- and background strain-matched. Animals were housed and bred in pathogen-free conditions in the vivarium at Thomas Jefferson University and maintained on a 12h:12h light/dark cycle. All experimental procedures were approved by the Thomas Jefferson University Institutional Animal Care and Use Committee (Protocol 01892).

Compound heterozygous mice (ACE2/G2019SLRRK2 ki) were generated by mating homozygous B6.Cg-Tg(K18-ACE2)2Prlmn/J (ACE2) with homozygous C57BL/6- Lrrk2tm4.1Arte (LRRK2^ki/ki^). In our hands, we empirically determined that there was no difference in neuropathological PD phenotype (loss of SNpc DA neurons) between heterozygous and homozygous LRRK2 mice, following the administration of a viral challenge. Genotypes of all animals used in this study were confirmed by PCR (Transnetyx, Cordova, TN).

### SARS-CoV-2 infection

All work involving the use of SARS-CoV-2 (USA-WA1/2020 (B1.1.1.7) and omicron B.1.1.529) were performed at BSL-3 biosafety conditions at Thomas Jefferson University. All viral studies and protocols were approved by Thomas Jefferson University’s Department of Environmental Health and Safety.

Prior to administration of virus, mice were anesthetized with 3% isoflurane. 25ul of various titers of WA-1/2020 or omicron in saline were intranasally administered (12.5ul in each nostril). Control mice were subjected to a sham procedure where they were similarly anesthetized and intranasally administered 25ul of sterile saline (Fig 1a). Following infection (or sham), animals were weighed daily and monitored for signs such as lack of grooming, hunched posture and reduced mobility for ten days post infection. Any mice exhibiting a loss of greater than 20% of the initial body weight were humanely euthanized. Additionally, if any animal showed any sign of intracerebral hemorrhage, either grossly or upon histological analysis these animals were not used for stererological assessment of SNpc DA neurons, Iba-1+ cell counts or cytokine analysis.

**Figure 1:**
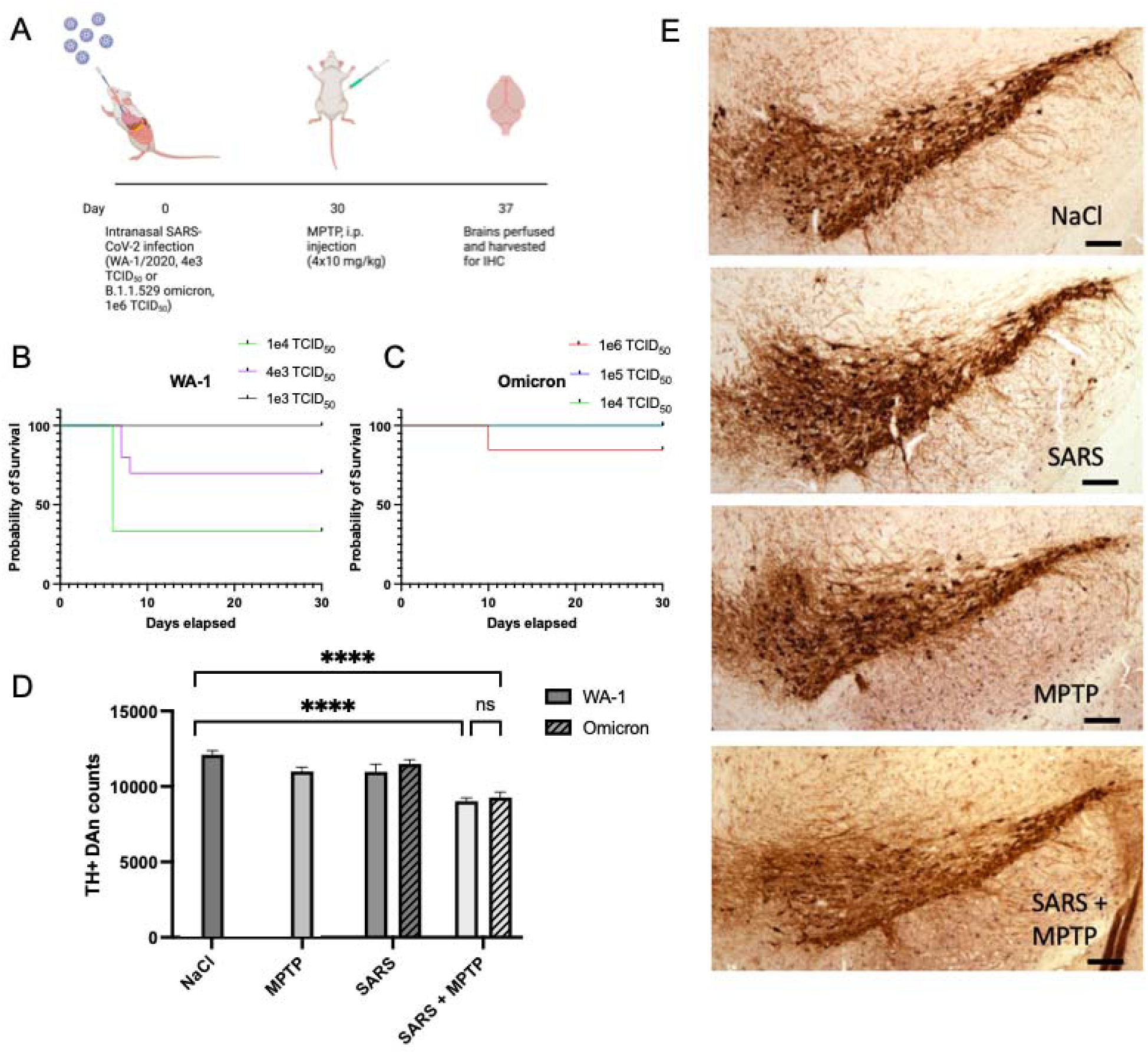
The omicron strain of SARS-CoV-2 induces a similar degree of neurodegeneration as WA-1 strain in an MPTP model of PD. (A) Schematic of mice used for analysis. (B) Kaplan-Meier survival curve of mice infected with three different titers of WA-1 strain of SARS-CoV-2. 1e3 TCID_50_: n=9; 4e3 TCID_50_: n=10; 1e4 TCID_50_: n=3. (C) Kaplan-Meier survival curve of mice infected with three different titers of WA-1 strain of SARS-CoV-2. 1e4 TCID_50_: n=3; 1e5 TCID_50_: n=3; 1e6 TCID_50_: n=13. (D) Quantification of TH+ DA neurons in the SNpc of k18-hACE2 mice revealing neurodegeneration in the SARS+MPTP groups. Data are mean ± SEM, n=4-9. ****p<0.0001. (E) Representative images of TH+ DA neurons in the SNpc of mice treated with NaCl, SARS alone (omicron strain), MPTP alone, and SARS(omicron)+MPTP. Scale bar=100um.

### MPTP and Paraquat administration

A subpathogenic dose (40 mg/kg)^60^ of 1-methyl-4-phenyl-1,2,3,6-tetrahydropyridine (MPTP) was administered to mice using a schedule of four i.p. injections of 10 mg/kg MPTP (Biosynth, Staad, Switzerland; cat #IM57898), each separated by 2 hours. Paraquat (PQ, Sigma Aldrich, St Louis, MO; cat #36541) was administered as three 20 mg/kg i.p. injections (total of 60 mg/kg total), each separated by 7 days. Control mice in each experiment were administered an equivalent volume of sterile saline. Seven days following the last toxin injection, mice were anesthetized with Avertin and following loss of deep tendon and corneal reflexes, were perfused with 5 ml 1x PBS followed by 25-30 ml 3% paraformaldehyde in 1X PBS.

### Immunization with SARS-CoV-2 vaccines

Mice were anesthetized with 3% isoflurane, and then injected intramuscularly (IM) with 1ug S-2P mRNA-LNP (obtained from Dr. Norbert Pardi, University of Pennsylvania, Philadelphia PA) diluted to a final volume of 100ul (50ul in each hind leg). Control animals were injected with the same volume of luciferase mRNA-LNP^61^. To assess induction of an immune response, 200ul of blood was collected through a puncture of the retroorbital vein prior to immunization as well as at day 21 following immunization. Mice were challenged with SARS- CoV-2 infection at 30 days following initial immunization.

For the CORAVAX vaccine (obtained from Dr. Mattias Schnell, Thomas Jefferson University, Philadelphia PA), mice were anesthetized with 3% isoflurane and injected IM with 10ug of vaccine diluted to a final volume of 100ul (50ul in each hind leg). 30 days following the priming dose, a booster dose (also 10ug) was administered IM. Control animals were injected with vehicle (sterile PBS). To assess induction of an immune response, 200ul of blood was collected through a puncture of the retroorbital vein prior to immunization, as well as at days 21, 25 and 56 following initial immunization. Mice were challenged with SARS-CoV-2 infection at 60 days following initial immunization.

### ELISA assay

An indirect enzyme linked immunosorbent assay (ELISA) was performed according to manufacturer instructions (AcroBiosystems, Newark, DE; cat #RAS-T045). Briefly, serum samples were diluted 1:100, and incubated at 37°C in a microplate with immobilized SARS- CoV-2 Spike (S1). Following washes, the plate was then incubated at 37°C with HRP-conjugated secondary antibody. Following washes, substrate solution was added, and the plate was incubated at 37°C. The reaction was stopped after 20 mins with the addition of a stop solution, and intensity of absorbance was measured at 450nm and 630nm. OD_450 nm_-OD_630 nm_ was used to reduce background noise. A standard curve was generated with a dilution series of provided anti-SARS-CoV-2 antibody (mouse IgG) and used to calculate antibody concentrations. All samples were run in duplicates.

### Protein extraction and Luminex assay

Substantia nigra (brain) and lung tissue was collected following exsanguination via transcardial perfusion with 1X PBS (pH 7.4) and stored at -80°C until they were assayed. Brain tissues were homogenized using a motorized pestle mixer (Cole Parmer, Vernon Hills, IL; cat #EW-44468-25) in Bio-Plex cell lysis buffer, containing Factor 1 and Factor 2 at manufacturer- recommender concentrations (Bio-Rad, Hercules, CA; cat #171304011). Lung tissues were similarly homogenized in RIPA buffer (Thermo Fisher, Waltham, MA; cat #89900) containing protease inhibitor (Thermo Fisher, Waltham, MA; cat #78429) and phosphatase inhibitor (Thermo Fisher, Waltham, MA; cat #78420) at manufacturer-recommended concentrations.

Blood was collected by cardiac puncture and transferred into EDTA-treated tubes. Serum was separated through centrifugation at 2000g and stored at -20°C. The total protein concentration of each sample was determined using a DC protein assay (Bio-Rad, Hercules, CA; cat #5000111).

Substantia nigra, lung and serum samples were prepared for cytokine analysis in 96-well plates using Milliplex Map kit (Millipore, Burlington, MA; cat #MCYTOMAG-70K), as previously described^62^. The plates were run on a Luminex200^TM^ platform, and the data was analyzed using Belysa^®^ software. All samples were run in duplicates.

### Immunohistochemistry

Mice were transcardially perfused with 1x PBS (pH 7.4), followed by 3% paraformaldehyde (PFA) in PBS (pH 7.4). Brains were dissected out of the skull and post-fixed in 3% PFA overnight at 4°C. For preparation of sections, brains were dehydrated through a graded series of ethanols and xylenes, and embedded and blocked in paraffin. 10um serial sections were cut in the coronal plane and mounted onto Unifrost charged slides (Azer Scientific, Morgantown, PA; cat #EMS200+). Every other slide was dual-labeled for tyrosine hydroxylase (TH) (1:250, Sigma-Aldrich, St Louis, MO; cat #T2928) and ionized calcium-binding adaptor molecule 1 (Iba-1) (1:250, Fujifilm Wako, Richmond, VA; cat #019-19741), as described previously^34^.

### Stereologic Assessment of Cell Number

SNpc DA neurons (combined counts of TH^+^ and TH^-^, nissl^+^ DA neurons) and Iba-1+ microglia in the SNpc were stereologically-assessed using either the physical dissector (SNpc DA neurons) or optical fractionator (microglia) with Stereo Investigator (version 2019.1.3, MicroBrightField, Colchester, VT), as described previously^63^. SNpc DA neurons were counted if they were TH^+^ and had a clear and complete nucleus. To assure that any change in SNpc DA neuron number was due to cell death, sections were counterstained with a Nissl stain and examined for the presence of TH^-^, Nissl^+^ DA neurons. Iba-1+ microglia were classified as being in a “resting” or “activated” state depending on their morphology; resting microglia have small oval soma that average 3um or less in diameter, and long fine processes, whereas activated microglia have irregular soma greater than 3um in diameter, and short thickened processes^30^.

### Statistical methods

Differences between groups were analyzed by t-tests (for two group comparisons) or ANOVA (comparisons of more than two groups). a priori post-hoc analysis (Tukey) was performed if overall statistical significance was found with ANOVA. A p-value of <0.05 was considered significant. All statistical analyses were performed and visualized using GraphPad Prism^®^, version 10.1.1.

## Results

### The omicron strain of SARS-CoV-2 induces a similar degree of neurodegeneration as WA- 1/2020 strain in an MPTP model of PD

We previously reported an approximate loss of 27% of SNpc DA neurons in ACE2 mice infected with the WA-1/2020 strain of SARS-CoV-2^64^. Here, we investigated if a different strain of SARS-CoV-2 (omicron, strain B1.1.529) would induce the same sensitivity to MPTP as WA-1/2020. The B.1.1.529 (omicron) strain of the virus was chosen because of its reported difference in mortality and severity of symptoms compared to the WA-1/2020 strain in humans^54^.

As a first step, we examined morbidity and mortality of the WA-1/2020 strain (Fig 1b) and compared it to that of the omicron strain (B1.1.529) of SARS-CoV-2 (Fig 1c). Following inoculation with WA-1/2020, we observed a 67% mortality with 1e^3^ TCID_50,_ 27% mortality after 4e^3^ TCID_50_ and zero mortality after 1e^3^ TCID_50_ (Fig 1b, suppl Fig 1b). For examination of mortality following omicron inoculation, mice were infected with either 1e^4^, 1e^5^, or 1e^6^ TCID_50_ of the omicron strain (B1.1.529) of SARS-CoV-2. We empirically determined that the two lowest titers, 1e^4^ and 1e^5^ TCID_50,_ had zero mortality while animals infected with the highest titer (1e^6^) exhibited a 15% mortality (Fig 1c). The mortality index of these SARS-CoV-2 strains reflects a similar trend in humans where the WA-1/2020 strain resulted in a much more severe infection than omicron ^65^. Based on these empirical studies we chose to examine the 4e^3^ TCID_50_ titer for WA-1/2020 and 1e^6^ TCID_50_ titer for omicron since these mimicked moderate infection in humans^60^.

To examine synergistic effects of SARS-CoV-2 infection with chemicals linked to parkinsonism, ACE2 animals were infected with either the WA-1/2020 or omicron strain, allowed to recover for 30 days and then injected with a subtoxic dose of MPTP (Fig 1a). We found that infection with the omicron strain induced a similar degree of neurodegeneration (24% loss of SNpc DA neurons compared to control (p<0.001) as we observed following infection with WA-1/2020 (27% SNpc DA neurons loss (p<0.001 compared to control) (Fig 1d), despite the observation that omicron infection resulted in significantly lower mortality. Neither WA- 1/2020, omicron or MPTP alone induced a significant loss of SNpc DA neurons compared to controls (Fig 1d). We also investigated SNpc neurodegeneration with a “mild” WA-1/2020 infection (1e^3^ TCID_50_ titer) followed by MPTP exposure and found no significant SNpc DA neuron loss (Supp Fig 1c) suggesting that SARS-CoV-2-induced SNpc DA neuron sensitivity is titer-dependent.

### Omicron induces a similar degree of SNpc microglial activation as WA-1, but with an altered immunological profile

To determine if infection with WA-1/2020 or B.1.1.529 omicron induced microglial activation in the SNpc alone or following MPTP exposure, we infected ACE2 mice with SARS- CoV-2, with or without 10 mg/kg x 4 MPTP. In both experimental conditions we did not observe any significant change in the total number of microglia (Fig 2a). However, when we compared changes in resting and activated microglia individually, we observed a significant reduction in the number of SNpc resting microglia of the SARS-CoV-2 infected (both strains) + MPTP group compared to control (31% with WA-1/2020, p<0.0001; 35% with omicron, p<0.0001), as well as a significant decrease compared to the SARS-CoV-2-alone group (24% with WA-1/2020, p<0.05; 22% with omicron, p<0.05) (Fig 2a). We also observed a significant increase in the number of activated microglia in the SNpc of the SARS-CoV-2 + MPTP group compared to control (220% with WA-1/2020, p<0.0001; 228% with omicron, p<0.0001) as well as a significant decrease compared to the SARS-CoV-2-alone group (129% with WA-1/2020, p<0.0001; 42% with omicron, p<0.0001) or MPTP alone (44% with omicron, p<0.05). We also measured a significant increase in activated microglia (70%, p<0.05) in the SNpc of ACE2 mice given a “mild” infection (1e^3^ TCID_50_ titer) of WA-1 followed by MPTP (Supp Fig 1d), even though the inoculation regimen did not induce SNpc DA neuron degeneration (Supp Fig 1c).

**Figure 2:**
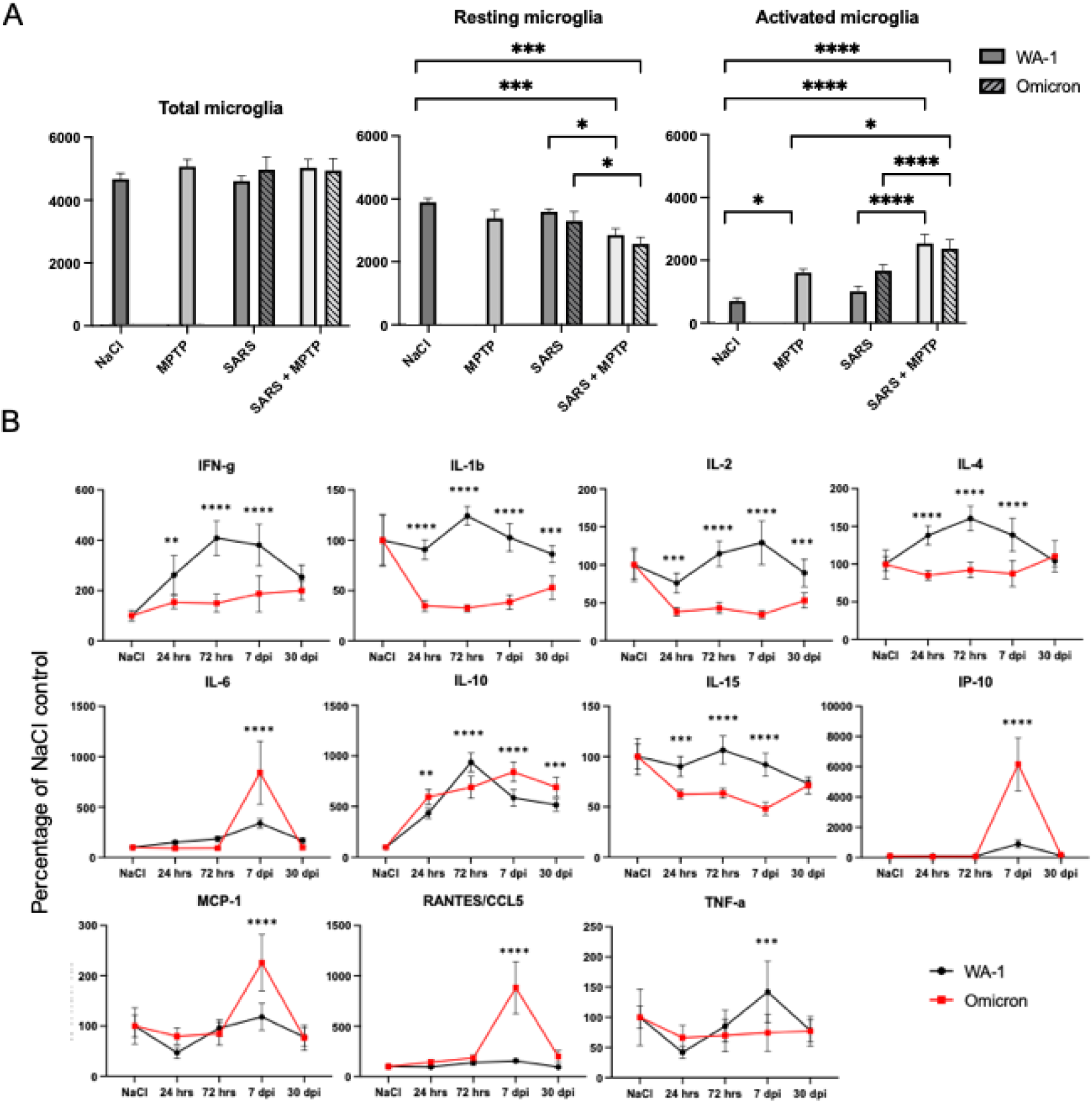
Omicron induces a similar degree of SNpc microglial activation as WA-1, but with an altered immunological profile. (A) Quantification of Iba-1+ total, resting, and activated microglia in the SNpc of k18-hACE2 mice revealing neuroinflammation in the SARS+MPTP groups. Data are mean ± SEM, n=4-13. *p<0.05, ***p<0.001, ****p<0.0001. (B) Cytokine and chemokine profiles in the SN comparing the time course of WA-1 (4e3 TCID_50_) and omicron (1e6 TCID_50_) infections from 24hrs post- infection to 30dpi. Data are mean ± SEM, n=6-9. **p<0.01, ***p<0.001, ****p<0.0001. Data in SARS-CoV-2 groups are expressed as the percentage of NaCl control.

To further explore the immunological profile of ACE2 mice following a SARS-CoV-2 infection, we infected the animals with either 4e^3^ TCID_50_ titer of the WA-1/2020 strain or 1e^6^ TCID_50_ titer of the omicron strain of the virus and assessed changes in pro-inflammatory cytokines (IFNγ, IL-1β, IL-2, IL-6, IL-15, TNFα), anti-inflammatory cytokines (IL-4, IL-10), and chemokines (IP-10, MCP-1, CCL5) at early infection timepoints (24hrs, 72 hrs), at peak infection timepoint (7dpi) and after the resolution of infection (30dpi) in the brain (SN) and the periphery (lungs). In the SN (Fig 2c), we found that WA-1/2020, compared to omicron, induced significantly higher levels of all the pro-inflammatory cytokines investigated except IL-6 as early as 24hrs post-infection. While the majority of the increased in pro-inflammatory cytokines resolved to baseline levels, IL-1β and IL-2 continue to be elevated through 30dpi. Among the anti-inflammatory cytokines we observed an increase in IL-4, at the early/peak stages of WA- 1/2020 infection but no change following infection with omicron. IL-10 levels were significantly increased following both WA-1/2020 and omicron and these levels stayed elevated through 30 days post-infection. We also observed a trend in chemokine expression (IP-10, MCP-1 and CCL5). Here we saw no induction at early stages of infection but after 7 dpi measured a significant increase in expression; however, this was only observed after omicron infection and not WA-1/2020. In the lungs, omicron induces an elevation in almost all the cytokines and chemokines investigated, except IFNγ (Supp Fig 2b) but we have incidentally observed more hemorrhaging present in the lungs after WA-1 infection (Supp Fig 2a).

### SARS-CoV-2 does not induce susceptibility in a paraquat model of PD

These studies, as well as previous work^60^, show that infection with the WA-1/2020 strain of SARS-CoV-2 synergizes with subtoxic dose of MPTP to induce basal ganglia pathology. Here, we examined if the WA-1/2020 strain of SARS-CoV-2 could synergize with another toxin linked to parkinsonism, paraquat, to induce neuropathology in the SNpc. Paraquat (PQ) is an herbicide, thought to induce free radical generation through the lipid peroxidation of membranes^66^, which has been epidemiologically linked with a higher incidence of PD in communities that report long-term exposures ^11^. While PQ has been shown to induce PD phenotypes in some models^10,67^, we have consistently seen no significant SNpc neurodegeneration or neuroinflammation ^68^, which we have replicated in this study. In C57BL/6J mice, PQ did not cause any change in SNpc DA neurons (Fig 3a) or in the number of activated, resting and total microglia (Fig 3b) of mice treated with PQ compared to saline-treated controls. We then examined if PQ effected changes subsequent to infection with SARS-CoV-2 (WA- 1/2020 strain) in ACE2 mice. Here, we did not observe any significant loss of SNpc DA neurons (Fig 3c) nor did we see any significant changes in numbers of activated, resting or total microglia (Fig 3d). This suggests that the interactions between viral infection and environmental agents linked to PD are toxin- or are, perhaps, mechanistically specific.

**Figure 3:**
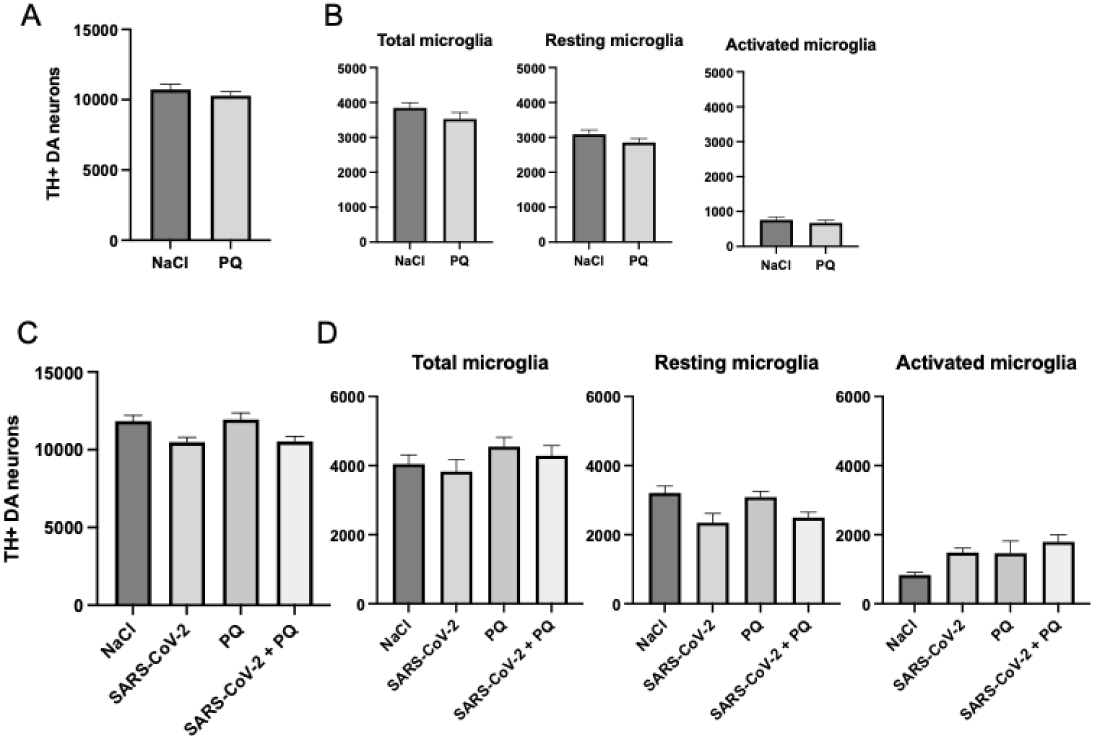
SARS-CoV-2 does not induce susceptibility in a paraquat model of PD. (A) Quantification of TH+ DA neurons in the SNpc of C57BL/6J mice revealing no neurodegeneration after treatment with PQ. Data are mean ± SEM, n=6-9. (B) Quantification of Iba-1+ total, resting, and activated microglia in the SNpc of C57BL/6J mice revealing no neuroinflammation after treatment with PQ. Data are mean ± SEM, n=6-9. (C) Quantification of TH+ DA neurons in the SNpc of k18-hACE2 mice revealing no neurodegeneration after treatment with SARS+PQ. Data are mean ± SEM, n=6-9. (D) Quantification of Iba-1+ total, resting, and activated microglia in the SNpc of k18-hACE2 mice revealing no neuroinflammation after treatment with SARS+PQ. Data are mean ± SEM, n=4-10.

### Mice carrying a G2019S LRRK2 mutation are more susceptible to WA-1 and omicron strains of SARS-CoV-2

Next, we investigated whether the presence of a familial PD mutation would increase the susceptibility to SARS-CoV-2-induced PD pathology. Here, compound heterozygous ACE2/G2019S LRRK2 ki mice were infected with 4e^3^ TCID_50_ titer of the WA-1/2020 strain or 1e6 of the B1.1.7 omicron strains of SARS-CoV-2 (Fig 4a). In the WA-1/2020-infected animals, we observed a 40% mortality rate compared to a 20% mortality rate when infected with 1e^6^ TCID_50_ titer of the omicron strain of the virus (Fig 4b), reflecting a trend seen not only with ACE2 mice (Fig 1b, c), but also outcomes of human patients^54^. Despite this difference in mortality and morbidity, we measured no difference in the loss of SNpc DA neurons at 30dpi between the two strains of the virus (Fig 4c). Both WA-1 and omicron infections in mice carrying a G2019S ki LRRK2 mutation led to a loss of approximately 20% of SNpc DA neurons (Fig 4c). However, the two strains of virus showed different patterns of microglial activation, with the omicron strain inducing a significantly higher number of total microglia (p<0.01) and resting microglia (p<0.001) compared to the WA-1/2020 strain (Fig 4d). Both strains induced the presence of a significant number of activated microglia compared to controls (WA-1/2020, p<0.05; omicron, p<0.001) (Fig 4d).

**Figure 4:**
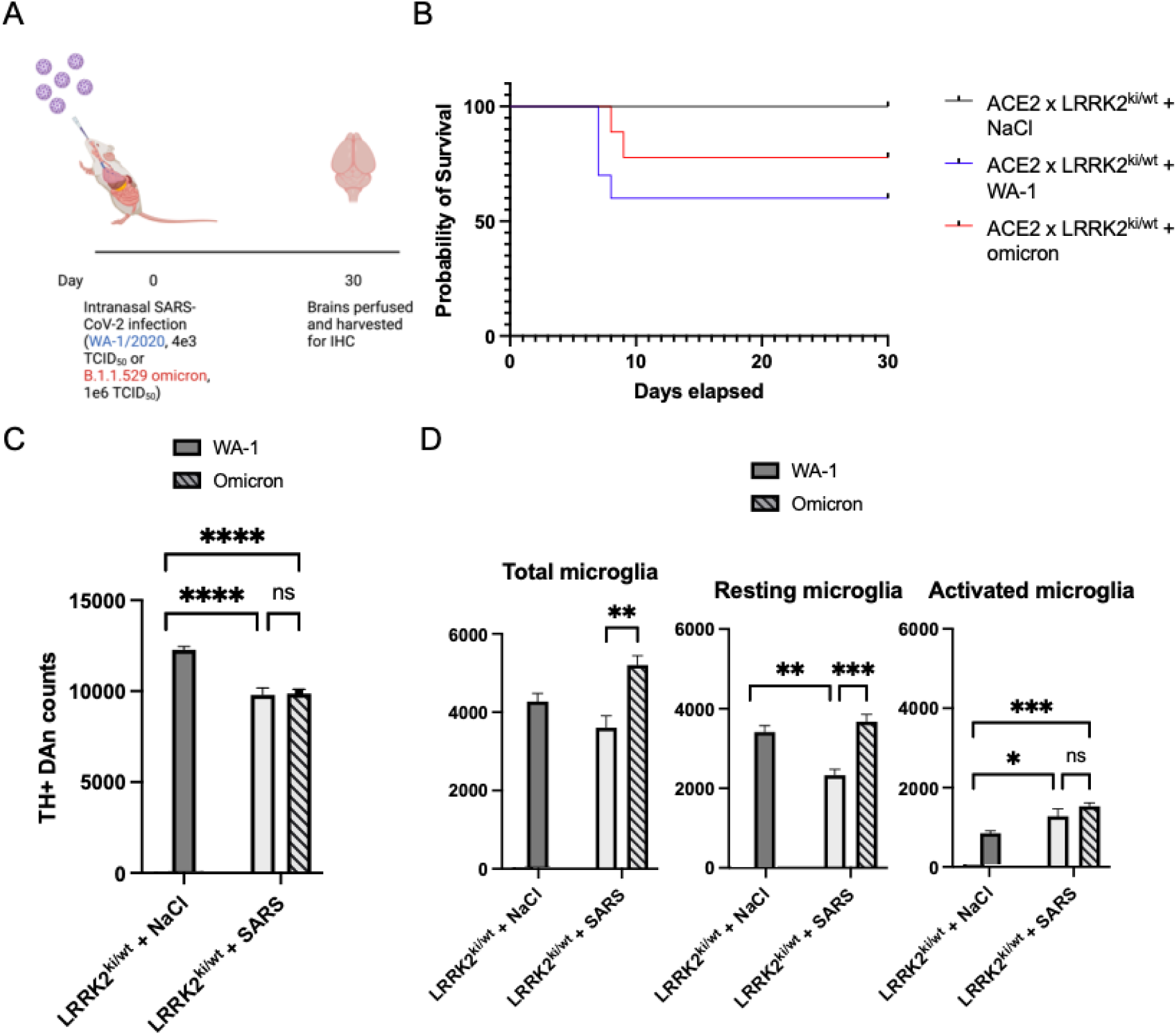
Mice carrying a LRRK2 mutation are susceptible to WA-1 and omicron strains of SARS-CoV-2. (A) Schematic of mice used for analysis. (B) Kaplan-Meier survival curve of ACE2/LRRK2 mice infected with WA-1 (4e3 TCID_50_) (n=10) or omicron (n=18), compared to sham controls (n=10). (C) Quantification of TH+ DA neurons in the SNpc of ACE2/LRRK2 mice revealing neurodegeneration in the SARS groups. Data are mean ± SEM, n=6-11. ****p<0.0001. (D) Quantification of Iba-1+ total, resting, and activated microglia in the SNpc of ACE2/LRRK2 mice revealing neuroinflammation in the SARS groups. Data are mean ± SEM, n=6-10. *p<0.05, **p<0.01, ***p<0.001, ****p<0.0001.

### ACE2/G2019S LRRK2 ki mice exhibit altered patterns of immunological changes after exposure to WA-1/2020 and omicron strains of SARS-CoV-2

To further investigate the interaction between SARS-CoV-2 infection and G2019S LRRK2, we examined the immunological profile of ACE2/G2019S LRRK2 ki compound heterozygous mice following a SARS-CoV-2 infection. We infected the animals with 4e^3^ TCID_50_ titer of the WA-1/2020 strain and then assessed changes in pro-inflammatory cytokines (IFNγ, IL-1β, IL-2, IL-6, IL-15, TNFα), anti-inflammatory cytokines (IL-4, IL-10), and chemokines (IP-10, MCP-1, CCL5) early after infection (24hrs, 72 hrs), at peak infection (7dpi) and after the resolution of infection (30dpi) in both brain (SN) and periphery (sera and lungs) (Fig 5a).

**Figure 5:**
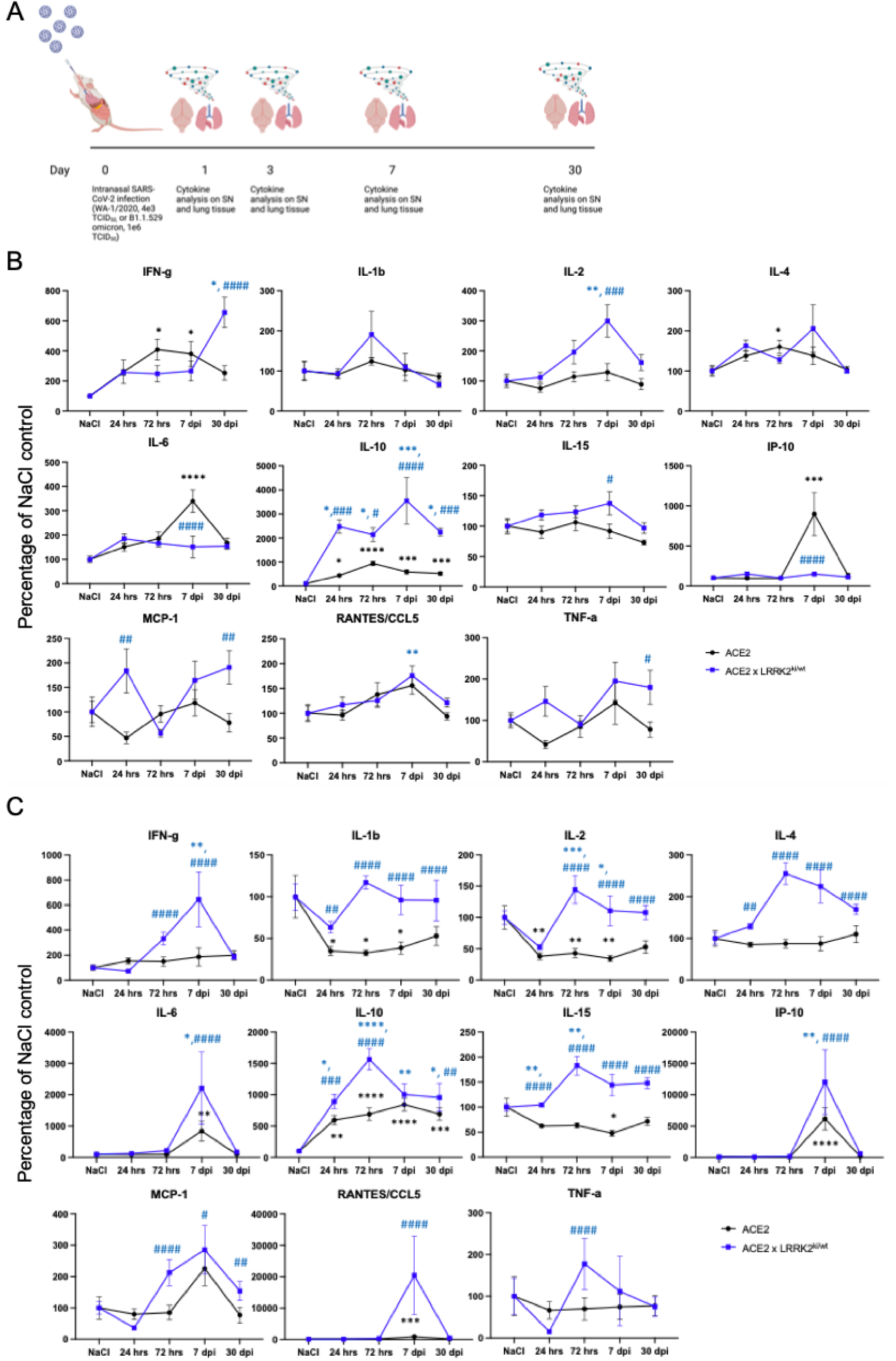
ACE2/LRRK2 mice exhibit altered patterns of immunological changes after exposure to WA-1 and omicron strains of SARS-CoV-2. (A) Schematic of mice used for analysis. (B) Cytokine and chemokine profiles in the SN of ACE2/LRRK2 and age-matched k18-hACE2 mice after infection with the WA-1 strain of SARS-CoV-2 (4e3 TCID_50_), looking at a time course from 24hrs post-infection to 30dpi. Data are mean ± SEM, n=6-9. **p<0.01, ***p<0.001, ****p<0.0001 compared to NaCl control, ^#^p<0.05, ^##^p<0.01, ^###^p<0.001, ^####^p<0.0001 compared to SARS-infected k18-hACE2 mice. For each genotype, data in SARS-CoV-2 groups are expressed as the percentage of NaCl control. (C) Cytokine and chemokine profiles in the SN of ACE2/LRRK2 and age-matched k18-hACE2 mice after infection with the omicron strain of SARS-CoV-2 (1e6 TCID_50_), looking at a time course from 24hrs post-infection to 30dpi. Data are mean ± SEM, n=6-9. **p<0.01, ***p<0.001, ****p<0.0001 compared to NaCl control, ^#^p<0.05, ^##^p<0.01, ^###^p<0.001, ^####^p<0.0001 compared to SARS-infected k18-hACE2 mice. For each genotype, data in SARS-CoV-2 groups are expressed as the percentage of NaCl control.

Following infection with WA-1/2020 SARS-CoV-2, ACE2/G2019S LRRK2 ki mice demonstrated a significantly higher increase in the level of the pro-inflammatory cytokines IFNγ, IL-2, IL-15, TNFα and chemoattractant factors MCP-1 and RANTES/CCL5 in the SN, with IFNγ, TNFα and MCP-1 remaining at increased levels compared to age-matched ACE2 controls through 30dpi (Fig 5b). However, following WA-1/2020 infection we also detected an increase in the anti-inflammatory cytokine IL-10 as early as 24 hrs post-infection which remained elevated through 30dpi (Fig 5b). While we see similar increases in pro-inflammatory cytokines (IFNγ, IL-1β, IL-6, IL-15) and chemokines (IP-10, MCP-1, CCL5) in the lungs, the inflammation in this tissue peaks at 7dpi and resolves by 30dpi (Supp Fig 3a). There is also a significant decrease in the anti-inflammatory cytokine IL-4, suggesting the virus can downregulate the anti-inflammatory mechanism in the lungs of ACE2/G2019S LRRK2 ki animals (Supp Fig 3a), but not in the brain (Fig 5b). Interestingly, we find negligible release of all investigated cytokines/chemokines in the sera of both ACE2 (Supp Fig 4a) and ACE2/G2019S LRRK2 ki (Supp Fig 4b) animals when infected with WA-1/2020, compared to the cytokine/chemokine response in the lungs and SN. This suggests that the infection as well as Fig 5. subsequent inflammation in the periphery is limited to the lungs in these mice, and unlike bacterial infections that model sepsis^62,69^, there is no spillover into the bloodstream.

To investigate the immunological profile of ACE2/G2019S LRRK2 ki infection with the omicron strain, we infected mice with 1e^6^ TCID_50_ titer of the virus and assessed the same analytes and timepoints as with WA-1 in the brain (SN) and periphery (lungs). Here, we see significant increases in all the proinflammatory cytokines and chemokines measured from the SN of ACE2/G2019S LRRK2 ki mice compared with their age-matched ACE2 controls with IL-1β, IL-2, IL-15 and MCP-1 remaining upregulated through 30dpi (Fig 5c). We also see a significant concurrent increase in anti-inflammatory cytokines IL-4 and IL-10 starting at 24hrs post-infection, which remain elevated through 30dpi (Fig 5c).

While changes were seen in the brain of ACE2/G2019S LRRK2 ki mice following infection with both strains of SARS-CoV-2, we did not see a similar increase in the proinflammatory cytokines or chemokines in the lungs except for upregulation of IFNγ and RANTES/CCL5 (Supp Fig 3b). There is also a significant downregulation of the anti-inflammatory cytokine IL-10 starting at 72 hours post-infection through 30dpi, again suggesting that the omicron strain is also able to downregulate the anti-inflammatory response in the periphery. The differences we see in the expression of proinflammatory cytokines between WA- 1/2020 and omicron in the lungs might explain the difference in mortality in ACE2/G2019S LRRK2 ki mice (Fig 4b), since lung damage and pneumonia play a large role in mortality from SARS-CoV-2^70^. Conversely, the lack of difference in the immunological pattern in the SN seen after both the WA-1/2020 and omicron strains may explain the similarities in the observed neuropathological changes induced by both strain of SARS-CoV-2 administered to ACE2/G2019S LRRK2 ki mice (Fig 4c).

### Immunization with S-2P mRNA-LNP, an mRNA vaccine against SARS-CoV-2, confers protection against neurodegeneration and neuroinflammation in the SNpc of k18-hACE2 mice, but not in mice carrying a G2019S LRRK2 mutation

mRNA-based vaccines have emerged as a novel modality in the fight against SARS- CoV-2, and make up the majority of immunizations given in the USA^71^. Here, we examined if a vaccine targeting the spike (S) surface glycoprotein that mediates viral cell entry and binding to the ACE2 receptor could protect mice from the synergy observed from the interactions of SARS- CoV-2 and the mitochondrial toxin MPTP. The S-2P vaccine is designed with a modified Spike protein containing K986P and V987P substitutions that stabilize S in the prefusion conformation that is attached to a lipid nanoparticle (LNP) for increased mRNA stability^72^. A single dose of this vaccine has been shown to elicit robust immune cell and neutralizing antibody responses.^73^ In this study, we investigated whether this protection extends to the SARS-mediated neurodegeneration and neuroinflammation we have described previously.

First, we performed a dose-finding study to look at the efficacy of the S-2P vaccine in rescuing parkinsonian phenotypes at 1ug and 5ug. Mice were given a 1 ug or 5 ug single dose of the S-2P vaccine, and then 30 days later were challenged with SARS-CoV-2 (WA-1) infection followed after another 30 days by subtoxic MPTP (Fig 6a). 7 days after the MPTP was administered, stereological examination of the SNpc DA neurons found no difference between the 1ug and 5ug dose in terms of vaccine-mediated rescue of neurodegeneration (Supp Fig 5a). In addition, we found rescue of neuroinflammation in the SARS+MPTP group with both doses of the vaccine (Supp Fig 5b). Thus, all subsequent experiments were carried out with the 1ug dose of the S-2P mRNA-LNP vaccine.

**Figure 6:**
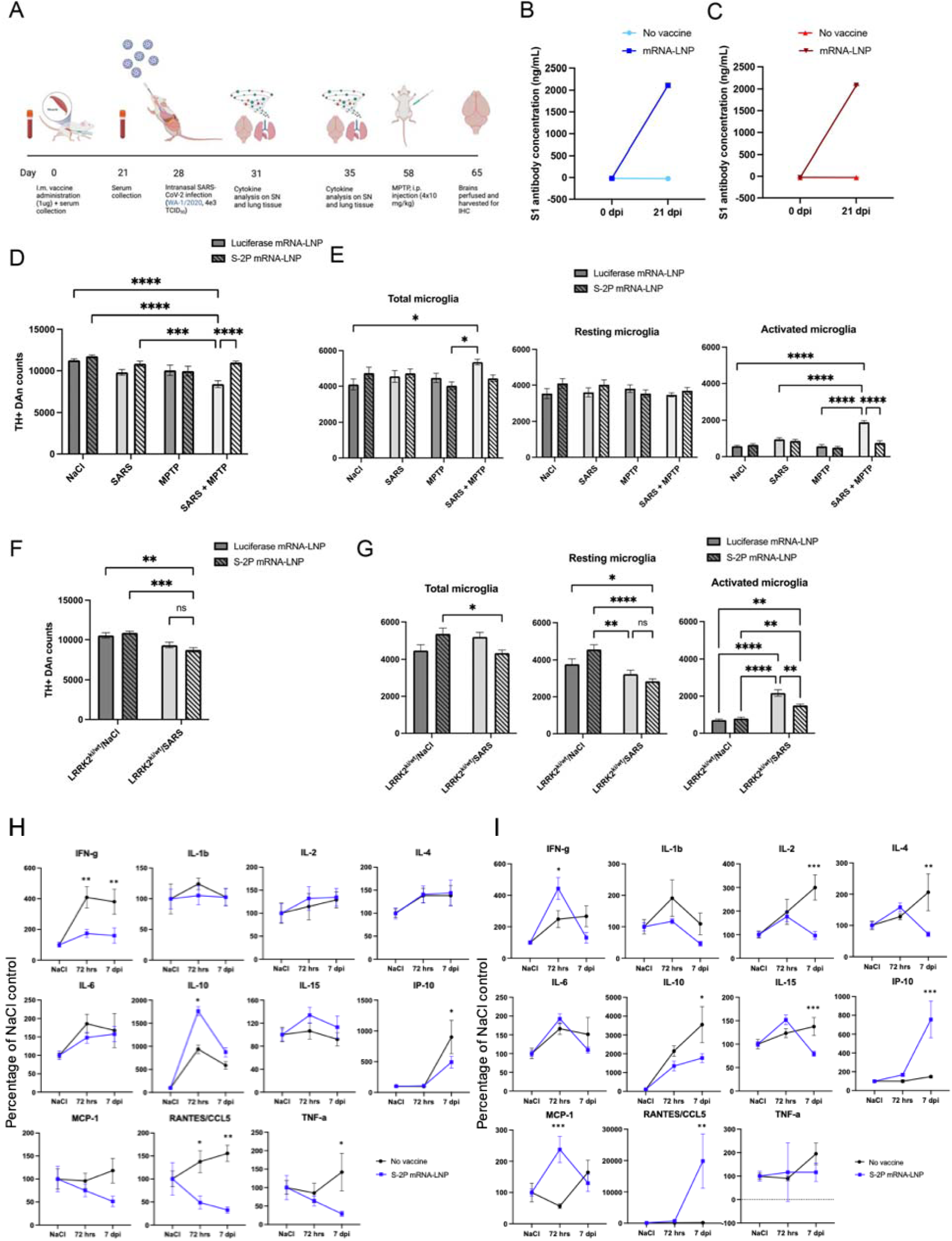
Immunization with S-2P mRNA-LNP, an mRNA vaccine against SARS-CoV-2, confers protection against neurodegeneration and neuroinflammation in the SNpc of k18- hACE2 mice, but not in mice carrying a LRRK2 mutation. (A) Schematic of mice used for analysis. (B) IgG antibody concentrations of SARS-CoV-2 spike (S1) protein in k18-hACE2 mice after immunization with S-2P mRNA-LNP. (C) IgG antibody concentrations of SARS-CoV-2 spike (S1) protein in ACE2/LRRK2 mice after immunization with S-2P mRNA-LNP. (D) Quantification of TH+ DA neurons in the SNpc of k18-hACE2 mice revealing rescue of neurodegeneration in the SARS+MPTP groups when previously immunized with S-2P mRNA-LNP. Data are mean ± SEM, n=4-8. ***p<0.001, ****p<0.0001. (E) Quantification of Iba-1+ total, resting, and activated microglia in the SNpc of k18-hACE2 mice revealing rescue of neuroinflammation in the SARS+MPTP group when previously immunized with S-2P mRNA-LNP. Data are mean ± SEM, n=5-7. *p<0.05, **p<0.01, ***p<0.001, ****p<0.0001. (F) Quantification of TH+ DA neurons in the SNpc of ACE2/LRRK2 mice revealing failure to rescue neurodegeneration in the SARS group when previously immunized with S-2P mRNA-LNP. Data are mean ± SEM, n=4-7. **p<0.01, ***p<0.001. (G) Quantification of Iba-1+ total, resting, and activated microglia in the SNpc of ACE2/LRRK2 mice revealing failure to rescue neuroinflammation in the SARS group when previously immunized with S-2P mRNA-LNP. Data are mean ± SEM, n=4-7. *p<0.05, **p<0.01, ****p<0.0001. (H) Cytokine and chemokine profiles in the SN of vaccinated and unvaccinated k18-hACE2 mice after infection with the WA-1 strain of SARS-CoV-2 (4e3 TCID_50_), at 72hpi and 7dpi. Data are mean ± SEM, n=6-8. *p<0.05, **p<0.01. Data in SARS-CoV-2 groups are expressed as the percentage of NaCl control. (I) Cytokine and chemokine profiles in the SN of vaccinated and unvaccinated ACE2/LRRK2 mice after infection with the WA-1 strain of SARS- CoV-2 (4e3 TCID_50_), at 72hpi and 7dpi. Data are mean ± SEM, n=6-8. *p<0.05, **p<0.01, ***p<0.001. Data in SARS-CoV-2 groups are expressed as the percentage of NaCl control.

Both ACE2 (Fig 6b) and ACE2/G2019S LRRK2 ki mice (Fig 6c) immunized with 1ug S- 2P mRNA-LNP vaccine developed high levels of serum antibodies against SARS-CoV-2 spike (S1) protein (p<0.0001 compared to unvaccinated controls) as measured at 21days post-immunization. Mice were immunized with S-2P mRNA-LNP or control luciferase mRNA-LNP, followed by inoculation with SARS-CoV-2 and then MPTP 30 days post-infection. In the luciferase group, we observed a 25% loss of TH+ and TH-, Nissl + DA neurons (p<0.001) compared to mice administered intranasal saline; similar to our previously reported study^60^.

However, immunization with the S-2P vaccine completely rescued this neurodegeneration (Fig 6d). In the luciferase vaccine control group, we observed a 30% increase in the total number of microglia when treated with SARS+MPTP compared to saline (p<0.05). We also observed a 233% increase in activated microglia in the SARS+MPTP group, compared to saline (p<0.0001). This rise in inflammation was completely rescued when animals were immunized with the S-2P vaccine (p<0.0001) (Fig 6e). When we examined the S-2P vaccine effects in ACE2/G2019S LRRK2 ki mice, we did not observe protection against SARS-mediated neurodegeneration. In fact, S-2P immunized ACE2/G2019S LRRK2 ki mice had a slightly higher degree of SNpc DA neuron loss (∼19%) compared to control mice immunized with the control luciferase vaccine (∼12%) (Fig 6f). Regarding neuroinflammation, immunization with control luciferase vaccine in ACE2/G2019S LRRK2 ki mice led to 205% increase in activated microglia in SARS-infected mice compared to saline controls (p<0.0001). Immunization with the S-2P vaccine did confer some degree of protection against inflammation, but still led to a 100% increase in activated microglia in SARS-infected mice compared to saline controls (p<0.01). We also observed a 38% decrease in resting microglia in the S-2P group when infected with SARS (p<0.0001), as well as a slight decrease in total microglia (p<0.05) (Fig 6g).

We then examined the immunological profiles of ACE2 (Fig 6h) and ACE2/G2019S LRRK2 ki mice (Fig 6i) vaccinated with S-2P following a SARS-CoV-2 infection. We assessed changes in pro-inflammatory cytokines (IFNγ, IL-1β, IL-2, IL-6, IL-15, TNFα), anti-inflammatory cytokines (IL-4, IL-10), and chemokines (IP-10, MCP-1, CCL5) early after infection (72 hrs), and at peak infection (7dpi) in both brain (SN) and periphery (lungs). In the SN of ACE2 mice, we found that the S-2P vaccine significantly reduced the pro-inflammatory cytokines IFNγ and TNFα, as well as chemokines IP-10 and CCL5. Vaccination also significantly increased the anti-inflammatory cytokine IL-10, showing an overall trend towards reduced inflammation (Fig 6h). We observed similar trends in the lungs of vaccinated ACE2 mice, with significant decreases in pro-inflammatory cytokines IL-1β, IL-2, IL-15 and TNFα as well as chemokines IP-10 and MCP-1 (Supp Fig 6a). In the lungs of ACE2/G2019S LRRK2 ki mice, we see similar reduction in inflammation, with significant decreases of pro-inflammatory cytokines IFNγ, IL-2, IL-6, IL-15 and TNFα and chemokines IP-10, MCP-1 and CCL5. We also observe an increase in the anti-inflammatory cytokine IL-4. (Supp Fig 6b). However, in the SN of ACE2/G2019S LRRK2 ki mice, we see a more mixed profile. Pro-inflammatory cytokines IL- 2 and IL-15 are significantly decreased in the vaccinated mice; however, IFNγ is significantly increased at 72 hrs. Both anti-inflammatory cytokines IL-4 and IL-10 are significantly decreased upon vaccination. We also observed significant increases in chemokines IP-10, MCP-1 and CCL5 in the vaccinated mice (Fig 6i). These data suggest that while the S-2P vaccine confers peripheral protection against SARS-CoV-2, reducing inflammation in the lungs and rescuing against SARS-induced mortality, it is not able to regulate the pro-inflammatory immune profile in the SN of ACE2/G2019S LRRK2 ki mice, and consequently, is unable to rescue SARS- mediated nigral neurodegeneration.

### Immunization with CORAVAX, a rabies vector-based vaccine against SARS-CoV-2, confers protection against neurodegeneration and neuroinflammation in the SNpc

Given that several millions of people worldwide were immunized with vaccines that utilized protein adjuvant-based technology (e.g, the Oxford-AstraZeneca vaccine ChAdOx1)^74^, we next investigated whether another protein-based vaccine would prove to be equally effective against SARS-CoV-2-induced neurodegeneration and neuroinflammation in either the MPTP or LRRK2 models of experimental PD. CORAVAX utilizes the S1 domain of the SARS-CoV-2 spike protein in an inactivated rabies vector, and has been shown to produce high antigen-specific serum antibody titers, as well as long-lived immunity for up to 1 year^75–77^.

Mice were primed with a 10ug dose of CORAVAX and boosted with an equal dose 30 days later. After an additional 30 days, ACE2 or ACE2/G2019S LRRK2 ki mice were challenged with SARS-CoV-2 (WA-1/2020) infection (Fig 7a). Mice immunized with CORAVAX showed 100% survival, whereas ACE2 and ACE2/G2019S LRRK2 ki mice given PBS shams suffered about 20% and 25% mortality, respectively (Fig 7d). Both ACE2 and ACE2/G2019S LRRK2 ki mice immunized with CORAVAX developed high levels of serum antibodies against SARS-CoV-2 spike (S1) protein (p<0.0001 compared to unvaccinated controls), and these levels remained stable after the booster dose at 28 dpi, until 56 dpi (Fig 7b and c). In ACE2 mice infected with SARS-CoV-2, then challenged 30 days later with MPTP (10mg/kg x 4 doses spaced 2 hours apart) we measured an approximate 25% loss of TH+ and TH-, Nissl + DA neurons (p<0.001) compared to mice administered intranasal saline; similar to our previously reported results^60^. Following immunization with CORAVAX, we found complete rescue of this MPTP-induced neurodegeneration and inflammatory response (Fig 7e). In contrast to the S-2P mRNA-LNP vaccine, we found that ACE2/G2019S LRRK2 ki mice immunized with CORAVAX were completely protected against the SNpc DA neuron degeneration that was measured in sham vaccinated mice (Fig 7g, p<0.01). CORAVAX also conferred a rescue of the increased neuroinflammation induced by SARS-CoV-2 in the MPTP model (Fig 7f, p<0.001) as well as the G2019S LRRK2 model (Fig 7h, (p<0.01)).

**Figure 7:**
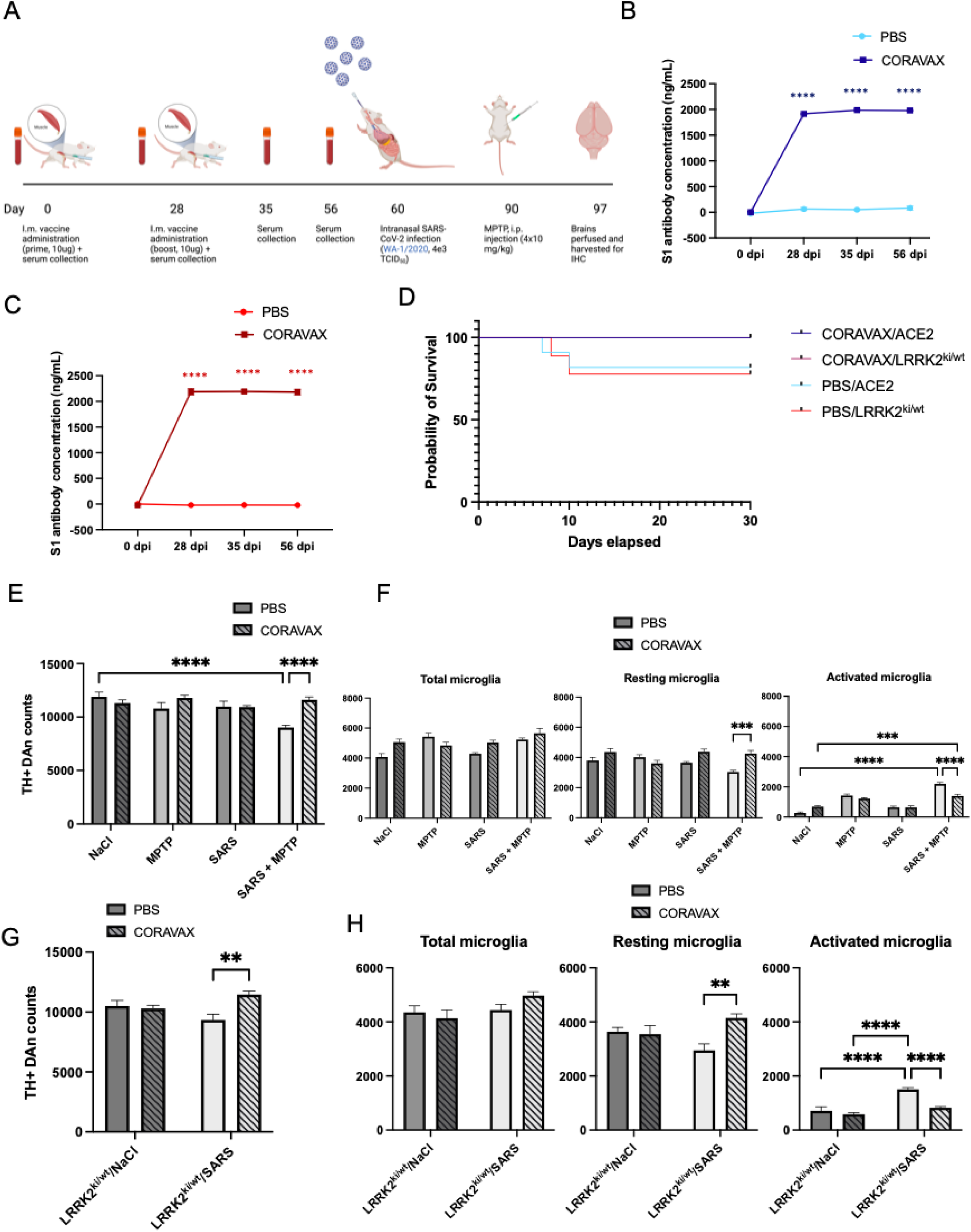
Immunization with CORAVAX, a rabies vector-based vaccine against SARS- CoV-2, confers protection against neurodegeneration and neuroinflammation in the SNpc. (A) Schematic of mice used for analysis. (B) IgG antibody concentrations against SARS-CoV-2 spike (S1) protein in k18-hACE2 mice after immunization with CORAVAX. (C) IgG antibody concentrations against SARS-CoV-2 spike (S1) protein in ACE2/LRRK2 mice after immunization with CORAVAX. (D) Kaplan-Meier survival curve of vaccinated (n=15) and unvaccinated (n=11) k18-hACE2 mice, and vaccinated (n=7) and unvaccinated (n=9) ACE2/LRRK2 mice infected with WA-1 at 4e3 TCID_50_ titer, revealing 100% survival in the vaccinated animals. (E) Quantification of TH+ DA neurons in the SNpc of k18-hACE2 mice revealing rescue of neurodegeneration in the SARS+MPTP groups when previously immunized with CORAVAX. Data are mean ± SEM, n=4-8. ****p<0.0001. (F) Quantification of Iba-1+ total, resting, and activated microglia in the SNpc of k18-hACE2 mice revealing rescue of neuroinflammation in the SARS+MPTP group when previously immunized with CORAVAX. Data are mean ± SEM, n=4-8. ***p<0.001, ****p<0.0001. (G) Quantification of TH+ DA neurons in the SNpc of ACE2/LRRK2 mice revealing rescue of neurodegeneration in the SARS group when previously immunized with CORAVAX. Data are mean ± SEM, n=4-7. **p<0.01. (H) Quantification of Iba-1+ total, resting, and activated microglia in the SNpc of ACE2/LRRK2 mice revealing rescue of neuroinflammation in the SARS group when previously immunized with CORAVAX. Data are mean ± SEM, n=4-7. **p<0.01, ****p<0.0001.

## Discussion

In this study, we demonstrate that infection with two different strains of the SARS-CoV-2 virus can induce peripheral and CNS inflammation along with increased risk for development of SNpc DA neuron loss secondary to some environmental toxins known to induce parkinsonism. We also show that carriers of the G2019S mutation in the LRRK2 gene have a dysregulated immune response to SARS-CoV-2 infection that is refractive to the prevention offered to wild-type mice.

Post encephalic basal ganglia disease has been reported as a sequalae to a number of different viral infections, including influenza, western equine encephalitic virus and West Nile Virus and SARS-CoV-2^27,29,32,78–81,64^. In this study, we expanded on these previous findings and now show that the omicron strain of SARS-CoV-2, despite causing much lower mortality and morbidity than the WA-1/2020 strain, induces a similar degree of cell loss of SNpc DA neurons and induced neuroinflammation secondary to MPTP. Regarding neuroinflammation, we also demonstrated that the secretory cytokine/chemokine profiles of mice infected with the WA- 1/2020 (B1.1.1.7) and omicron B1.1.529 strains of the virus were completely different, suggesting that each strain of SARS-CoV2 appears to be recognized differently by both the peripheral immune system as well as the intrinsic immune system of the brain. However, despite these differences in cytokine induction, we observed no difference in the SNpc DA neuron loss induced by WA-1/2020 or omicron, suggesting that perhaps there isn’t a single inflammatory driver that leads to this degeneration, but rather, a general threshold of inflammation that must be breached to induce these changes^82,83^. This hypothesis is also supported by our data that examined a low-titer, zero mortality infection with WA-1/2020, which still led to a small but significant increase in activated microglia – but no SNpc DA neuron degeneration.

To examine if any form of induced oxidative stress could synergize with the inflammation induced by SARS-CoV-2 infection, we examined whether paraquat dichloride, an herbicide epidemiologically linked with the development of Parkinson’s disease^9,11^, like MPTP could synergize with SARS-CoV-2 to induce SNpc DA neuron. As previously reported^68^, we did not measure any significant neurodegeneration or neuroinflammation induced by PQ alone nor when its administration followed SARS-CoV-2 infection. This dichotomy of effect suggests that the increased risk for development of neurodegeneration following SARS-CoV-2 infection is dependent on the specific type or mechanism underlying the “second hit”. Here, MPTP induces oxidative stress through the inhibition of mitochondrial complex I and IV^84^, while PQ is thought to act via direct lipid peroxidation of cell membranes^66^. A number of recent studies examining mitochondrial genome changes subsequent to SARS-CoV-2 infection have shown downregulation of a number of mitochondrial genes including Complex I^85–87^, which could provide an explanation why MPTP and not PQ induces this synergistic effect. An interesting future direction with relevance to public health could be to look at other mitochondrial toxins such as rotenone^88^, or through if other inducers of oxidative stress through lipid peroxidation such iron or manganese^89–91^ to examine if any of these preferentially increases – or not – secondary post-viral risk for the development of both experimental and clinical PD.

Aside from toxic and infectious exposures known to induce parkinsonism, several genes have been identified that either directly lead to development of PD or greatly increase the risk for its development^5–8^. In this study we examined interactions of COVID-19 infection with the G2019S mutation in the LRRK2 gene, one of the most common risk factor alleles contributing to familial PD^5,6^. LRRK2 is a ubiquitously expressed protein that is especially abundant in the brain, kidneys and lungs^92^. Additionally, it is also highly expressed in B-cells, T-cells, macrophages and monocytes of the immune system where functionally it contributes to phagocytosis and autophagy pathways^93^. When mutated at the G2019S locus, LRRK2 kinase activity is induced at abnormally high levels^92^ due to its own autophosphorylation^94^. G2019S LRRK2 carriers are known to have dysregulated immune profiles, with increased pro-inflammatory cytokines seen in the serum of even asymptomatic patients^95^. Given its high expression in lung tissue and immune cells, it is unsurprising that we saw higher mortality and immune dysregulation after infection with SARS-CoV-2 in ACE2 mice carrying this LRRK2 mutation compared to the mortality seen in ACE2 mice alone. In terms of this dysregulation, several cytokines and chemokines remained upregulated in the SN even at 30 dpi, suggesting that these mice exhibit a sustained secretory inflammatory response, which correlates with our observation that activated microglia remain elevated in the SN through this timeframe. We also found, similar to our MPTP-induced oxidative stress model, that both the WA-1/2020 (B1.1.1.7) and B.1.1.529 omicron strains of SARS-CoV-2 induced similar degrees of neurodegeneration and activated microglia in the SNpc of ACE2/G2019SLRRK2 ki mice despite differing cytokine profiles, suggesting that no one cytokine was responsible for this effect but that changes in many of these are needed to exceed a general threshold of increased inflammation to begin the pathological cascades that ultimately lead to SNpc DA neuron death. Incidentally, we observed a slightly higher mortality in female LRRK2 mutant mice compared to males when exposed to SARS-CoV-2 and that sex differences in the induced immune response might underlie this observation. A recent study has shown sex-dependent immune cell exhaustion in aged LRRK2 mutant mice^96^, with macrophages from female mice demonstrating hypophagocytosis and decreased antigen presentation, while macrophages from male mice needed a chronic inflammatory stimulus to observe the same phenotype.

In regard to inflammation, one of the features of a moderate to severe SARS-CoV-2 infection is the development of a peripheral “cytokine storm”^97^. Here, cellular entry of the virus into the lungs induces activation of Th1 cells which secrete proinflammatory factors including IL-2, IL-6, IL-10, TNFα and MCP-1^98^ as well as CD4 and CD8 T-cell lymphopenia^99^. Clinically, the prognosis and severity of patient infections has been shown to be directly correlated with the severity of the cytokine storm^100^. However, unlike clinical reports, we find negligible levels of any the cytokines/chemokines that we analyzed in sera of both ACE2 and ACE/G2019S LRRK2 ki compound heterozygotes. Although not in sera, we do see a peripheral immune response; but of the tissues we measured, this appears to be isolated to the lungs without spill over into the bloodstream. It is possible that this difference in human versus mouse is secondary to the titers we used to simulate a mild-moderate infection which were based on matching levels of mortality; and thus, moving to a higher viral titer might induce a peripheral cytokine storm in sera similar to those seen in humans. Alternately, we might have identified a caveat in rodent studies where there are clear species differences in response to infection. Whatever the case, in our hands we do not see any induced cytokine response in the blood, and therefore the mode of inflammatory transmission from the periphery to the brain remains unclear. It is also possible that the viral induction of inflammation in the brain is completely detached from systemic infection. This, however, would require direct viral infection of the nervous system. At the titers we examined, we see no evidence of viral neurotropism in the SN except in a few cases where we detected hemorrhage indicating a BBB breach. However, any animals with an indication of BBB disruption were not included in our analysis. It is also possible that direct signals to the brain arise from other levels of the nervous system including signals that emanate from the peripheral nervous system^101^ including the olfactory, trigeminal or vagus nerves^102–104^. It is also possible that circulating virus could communicate with the brain via interactions of cells present in the glymphatic system^105^. These hypothesis are supported by transcriptomic profiling of olfactory and brainstem tissues in post-mortem tissue of COVID patients^106^ as well the crosstalk between brain and periphery through glymphatic, GI and lymphatic systems^107,108^.

Finally, in this study, we examined the important question of whether SARS-CoV-2 vaccines can protect against the neurological phenotypes we have described. We examined this question using two different vaccine formulations: 1) the S-2P mRNA-based vaccine and 2) a protein-based vaccine built upon an inactivated rabies vector (CORAVAX). In wild-type ACE2 mice, we found that both these vaccines not only protect against mortality and morbidity, but they also protect the substantia nigra from SARS-CoV-2 mediated neuropathology. However, in the ACE2/G2019S LRRK2 ki mice, while immunization with CORAVAX rescued SNpc DA neuron loss and the activation of microglia, immunization with the S-2P mRNA vaccine did not. mRNA vaccinated ACE2/G2019S LRRK2 ki mice showed the same degree of loss and inflammation as the unvaccinated mice, as well as similar dysregulation of their SN cytokine profiles. Studies have shown differences in the production of specific IgG subtypes depending on the mechanism of the vaccine used – mRNA vaccines induce greater levels of IgG2, whereas protein vaccines induce more IgG1^109^. It is possible that this difference in antibody production, coupled with the already dysregulated immune system in LRRK2 mutants underlies their non-response to the mRNA-based vaccine. Alternatively, LRRK2 mutant animals may need a higher dose of vaccine than their wild type littermates to achieve the same degree of efficacy. Here, we used the same 1ug single dose in both our ACE2 and ACE2/G2019S LRRK2 ki mice. Further studies will be required to see whether a higher single dose or a booster dose of the mRNA vaccine before inoculation with SARS-CoV-2 would subsequently rescue the induced neurological phenotypes observed in mice carrying the G2019S mutation.

Despite the differences in vaccine response in WT or G2019S LRRK2 mice, these studies highlight the importance of timely and regular vaccinations against viral diseases like SARS- CoV-2, not only to protect against immediate mortality/morbidity from the infection, but also the protection of longer-term neurological health – although our study also raises the question of whether recommendations for specific types of vaccines are necessary for patients, depending on their genetic background. Additionally, a significant number of patients recovering from a COVID infection can present with persistent residual sequelae lasting greater than 3 months.

This Post-Acute Sequelae of SARS-CoV-2 Infection (PASC)^110^ or known colloquially as “long COVID”^111^ is a syndrome that manifests in a constellation of symptoms including fatigue, cognitive impairment, depression, and autonomic dysfunction; together or individually greatly impairing quality of life^112–114^. Recent studies have shown that patients with long COVID have significantly different circulating immune cell populations as well as chronic dysregulated inflammation^111,115,116^. Besides the potential for development of PASC, our study again^64^ implicates that prior infection with COVID-19, independent of viral strain, could be a trigger that increases the risk for developing PD, especially among the millions of people worldwide and carriers of the G2019S mutation who survived the COVID-19 pandemic prior to the development of vaccines and other therapeutics.

## Acknowledgements

The authors thank Dr. Norbert Pardi, University of Pennsylvania, for the generous gift of the S-2P mRNA-LNP vaccine and Dr. Matthias Schnell, Thomas Jefferson University for the gift of the rabies vector-based CORAVAX vaccine. We also thank Tabitha Rodriguez, Kallon Crowther, Matthew Byrne, Brian Montoya, Tytisha Harris and Thomas Kuret for assistance in the ABSL3 facility, and Elena Kozina for critical reading of the manuscript.

This study was funded by the NIH (NINDS R21 NS122280) and the Michael J Fox Foundation for Parkinson’s Research. The funders played no role in study design, data collection, analysis and interpretation of data, or the writing of this manuscript.

## Author Contributions

DC designed, performed experiments, analyzed data and wrote the paper. DK performed experiments and analyzed data. RJS designed, performed experiments, analyzed data and wrote the paper. All authors read and approved the final manuscript.

## Competing Interests

Authors DC, DK and RJS declare no financial interests. Author RJS serves as Associate Editor of this journal and had no role in the peer-review or decision to publish this manuscript.

## Data Availability

Data sharing is not applicable to this article as no datasets were generated or analyzed during the current study.

---

Supplementary Figure 1: Low titer SARS-CoV-2 infection does not induce susceptibility in an MPTP model of PD. (A) Schematic of mice used for analysis. (B) Kaplan-Meier survival curve of k18-hACE2 mice infected with WA-1 at 1e3 TCID_50_ titer (n=10), compared to sham controls (n=10). (C) Quantification of TH+ DA neurons in the SNpc of k18-hACE2 mice revealing no neurodegeneration in the SARS+MPTP groups. Data are mean ± SEM, n=4-6. (D) Quantification of Iba-1+ total, resting, and activated microglia in the SNpc of k18-hACE2 mice revealing slight neuroinflammation in the SARS+MPTP group. Data are mean ± SEM, n=4-6. *p<0.05.

Supplementary Figure 2: Gross and immune changes in the lungs of k18-hACE2 mice following infection with WA-1 and omicron strains of SARS-CoV-2. (A) Representative image of hemorrhaging present in the lungs of mice infected with WA-1 (top right), compared to NaCl (top left). Representative image of lungs infected with omicron (bottom). (B) Cytokine and chemokine profiles in the SN comparing the time course of WA-1 (4e3 TCID_50_) and omicron (1e6 TCID_50_) infections from 24hrs post-infection to 30dpi. Data are mean ± SEM, n=6-8. *p<0.05, **p<0.01, ***p<0.001, ****p<0.0001. Data in SARS-CoV-2 groups are expressed as the percentage of NaCl control.

Supplementary Figure 3: Altered lung cytokine/chemokine profiles in ACE2/LRRK2 mice following SARS-CoV-2 infection. (A) Cytokine and chemokine profiles in the lungs of ACE2/LRRK2 and age-matched k18- hACE2 mice after infection with the WA-1 strain of SARS-CoV-2 (4e3 TCID_50_), looking at a time course from 24hrs post-infection to 30dpi. Data are mean ± SEM, n=6-9. **p<0.01, ***p<0.001 compared to NaCl control, ^#^p<0.05, ^##^p<0.01, ^###^p<0.001 compared to SARS- infected k18-hACE2 mice. For each genotype, data in SARS-CoV-2 groups are expressed as the percentage of NaCl control. (C) Cytokine and chemokine profiles in the lungs of ACE2/LRRK2 and age-matched k18-hACE2 mice after infection with the omicron strain of SARS-CoV-2 (1e6 TCID_50_), looking at a time course from 24hrs post-infection to 30dpi. Data are mean ± SEM, n=5-9. **p<0.01, ***p<0.001, ****p<0.0001 compared to NaCl control, ^#^p<0.05, ^##^p<0.01, ^###^p<0.001, ^####^p<0.0001 compared to SARS-infected k18-hACE2 mice. For each genotype, data in SARS-CoV-2 groups are expressed as the percentage of NaCl control.

Supplementary Figure 4: Negligible release of cytokines/chemokines in the sera of ACE2 and ACE2/LRRK2 mice upon infection with WA-1/2020. (A) Cytokine and chemokine concentrations (reported as pg/mg of total protein) in the sera, lungs and SN of ACE2 mice after infection with WA-1 strain of SARS-CoV-2 (4e3 TCID_50_), looking at a time course from 24hrs post-infection to 30dpi. Data are mean ± SEM, n=6-10. (B) Cytokine and chemokine concentrations (reported as pg/mg of total protein) in the sera, lungs and SN of ACE2/LRRK2 mice after infection with WA-1 strain of SARS-CoV-2 (4e3 TCID_50_), looking at a time course from 24hrs post-infection to 30dpi. Data are mean ± SEM, n=5-10.

Supplementary Figure 5: 1ug and 5ug doses of the S-2P mRNA-LNP vaccine confer the same degree of neuroprotection in the MPTP model. (A) Quantification of TH+ DA neurons in the SNpc of k18-hACE2 mice revealing rescue of neurodegeneration in the SARS+MPTP groups when previously immunized with S-2P mRNA-LNP at both 1ug and 5ug dose. Data are mean ± SEM, n=4-8. ***p<0.001, ****p<0.0001. (B) Quantification of Iba-1+ total, resting, and activated microglia in the SNpc of k18-hACE2 mice revealing rescue of neuroinflammation in the SARS+MPTP group when previously immunized with S-2P mRNA-LNP at both 1ug and 5ug dose. Data are mean ± SEM, n=4-7. *p<0.05, ****p<0.0001.

Supplementary Figure 6: Vaccination with S-2P mRNA-LNP reduces lung inflammation following SARS-CoV-2 infection. (A) Cytokine and chemokine profiles in the lungs of vaccinated and unvaccinated k18-hACE2 mice after infection with the WA-1 strain of SARS-CoV-2 (4e3 TCID_50_), at 72hpi and 7dpi. Data are mean ± SEM, n=6-9. *p<0.05, **p<0.01. Data in SARS-CoV-2 groups are expressed as the percentage of NaCl control. (B) Cytokine and chemokine profiles in the lungs of vaccinated and unvaccinated ACE2/LRRK2 mice after infection with the WA-1 strain of SARS-CoV-2 (4e3 TCID_50_), at 72hpi and 7dpi. Data are mean ± SEM, n=5-7. *p<0.05, **p<0.01, ***p<0.001. Data in SARS-CoV-2 groups are expressed as the percentage of NaCl control.

## Notes

### Competing Interest Statement

The authors have declared no competing interest.

