## Supplementary figures and images for "Environmental exposures and familial background alter the induction of neuropathology and inflammation after SARS-CoV-2 infection"

### Suppl fig 1

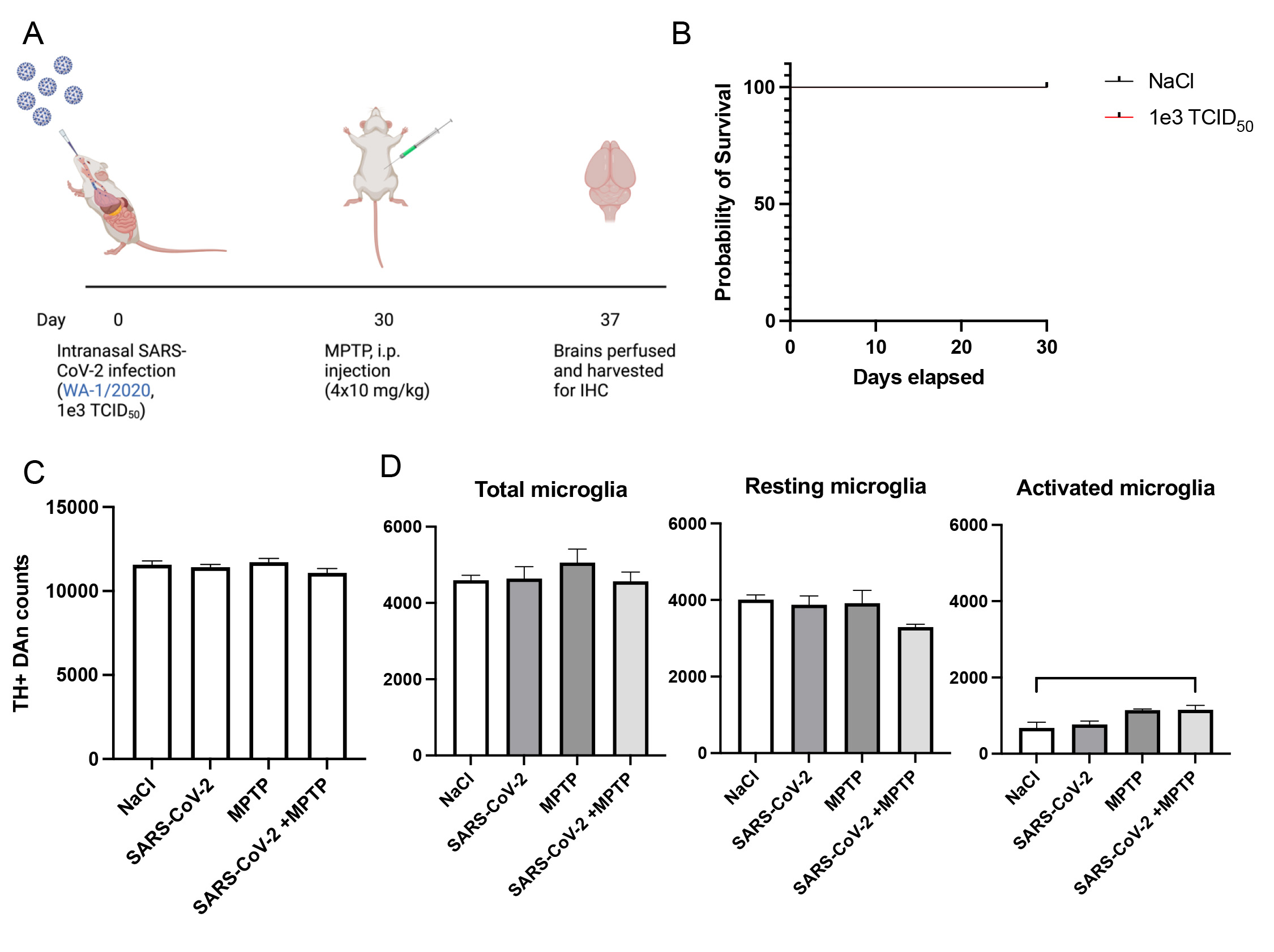

### Suppl fig 2

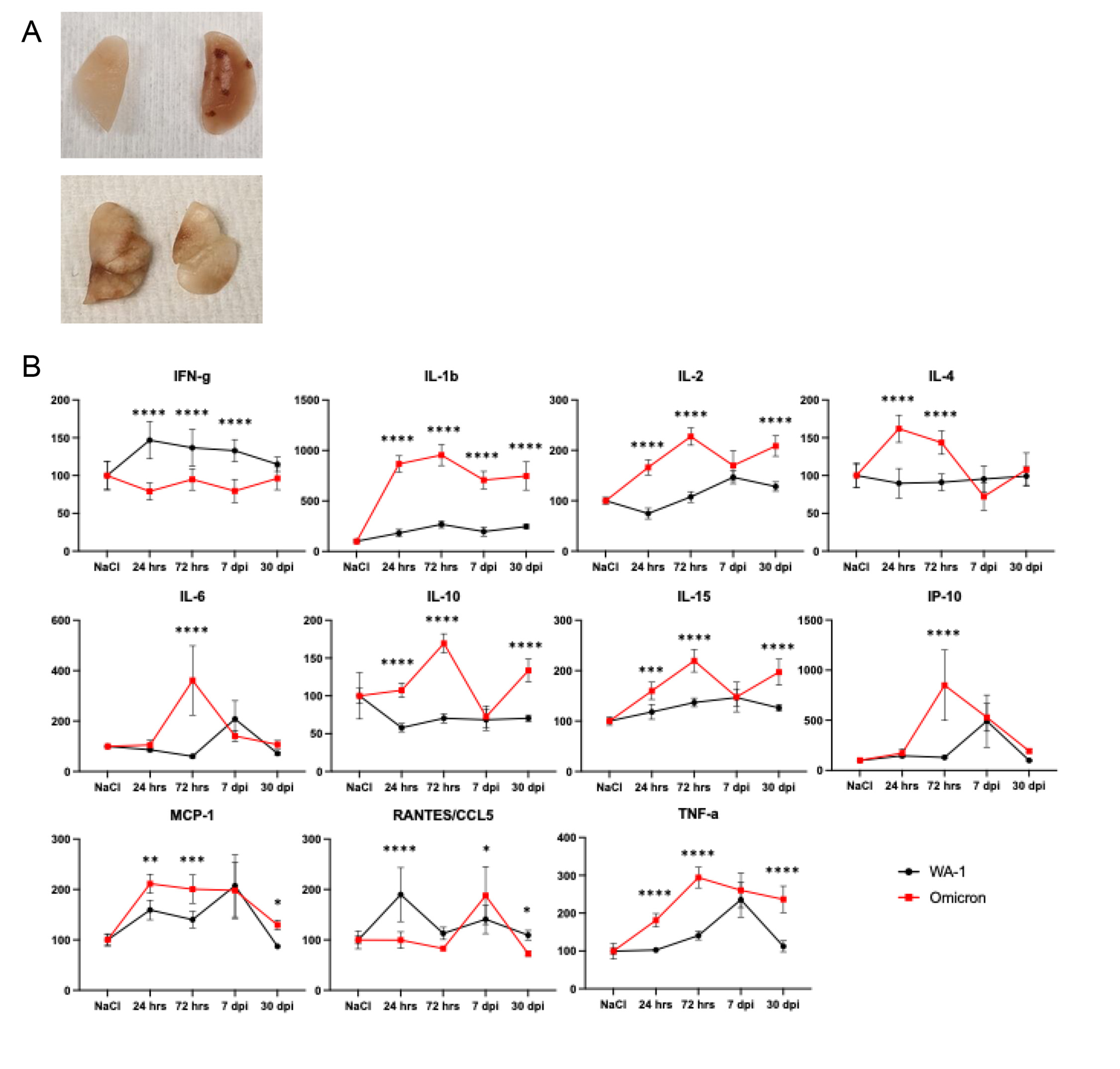

### Suppl fig 3

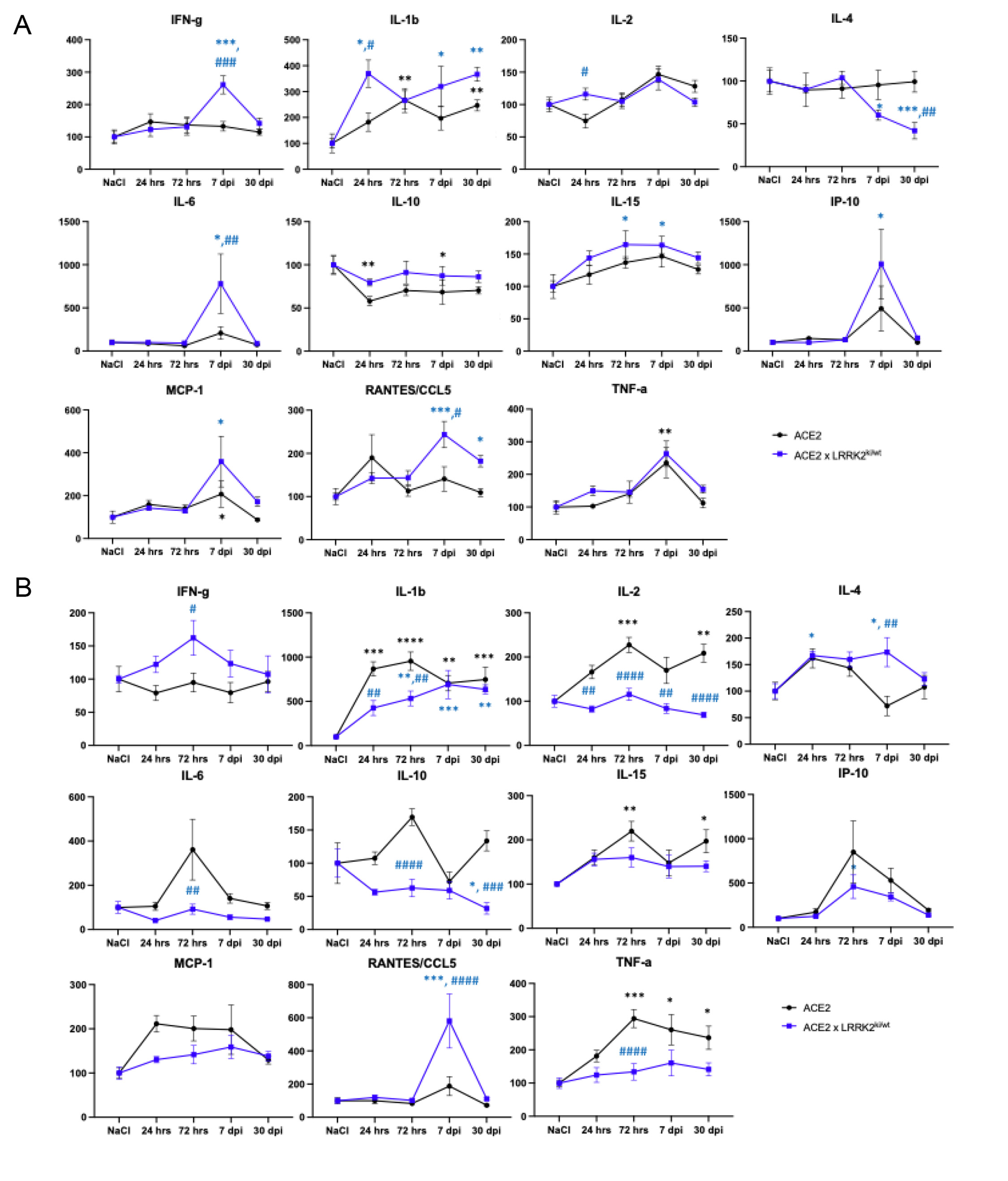

### suppl fig 4

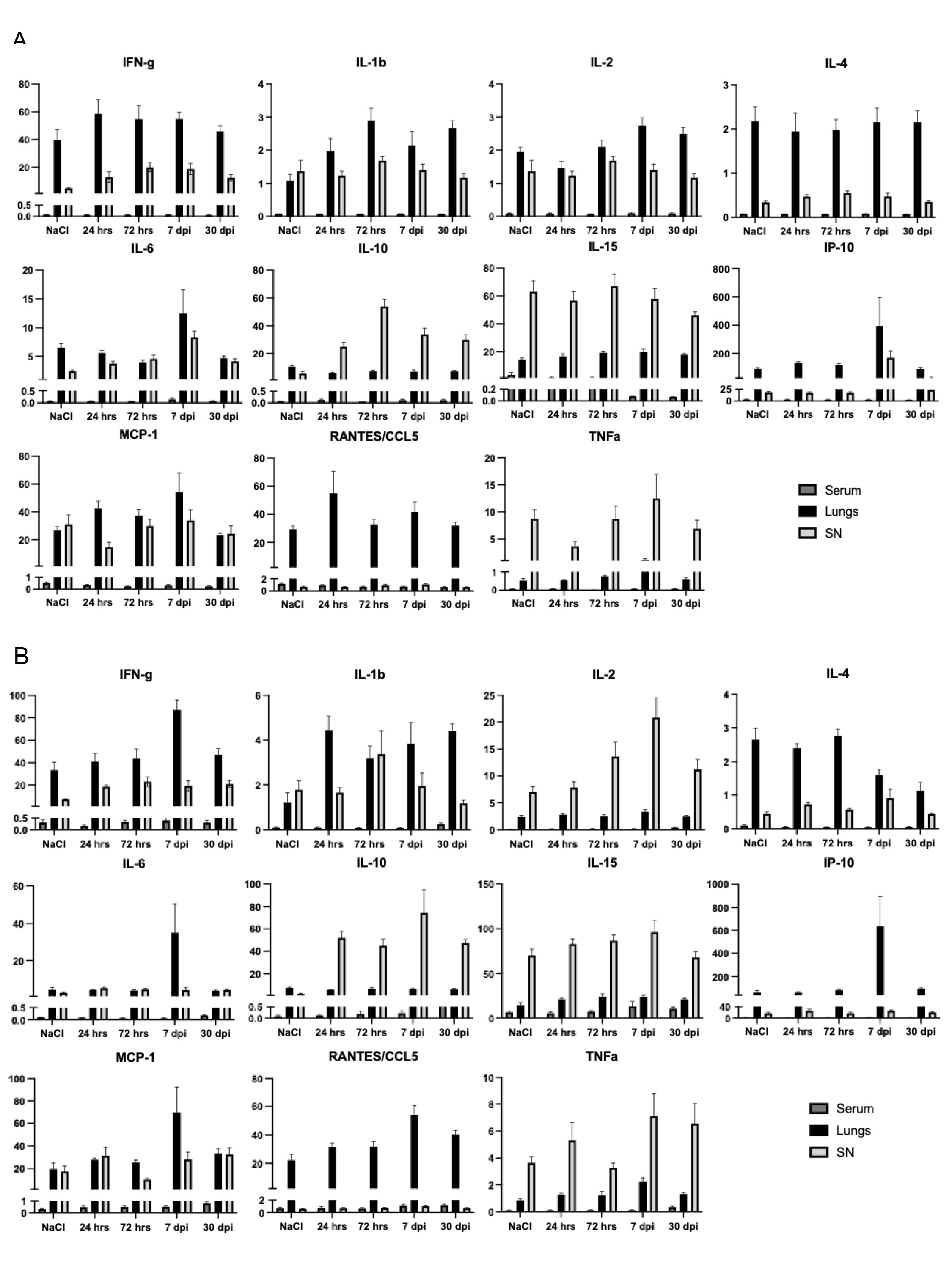

### Suppl fig 5

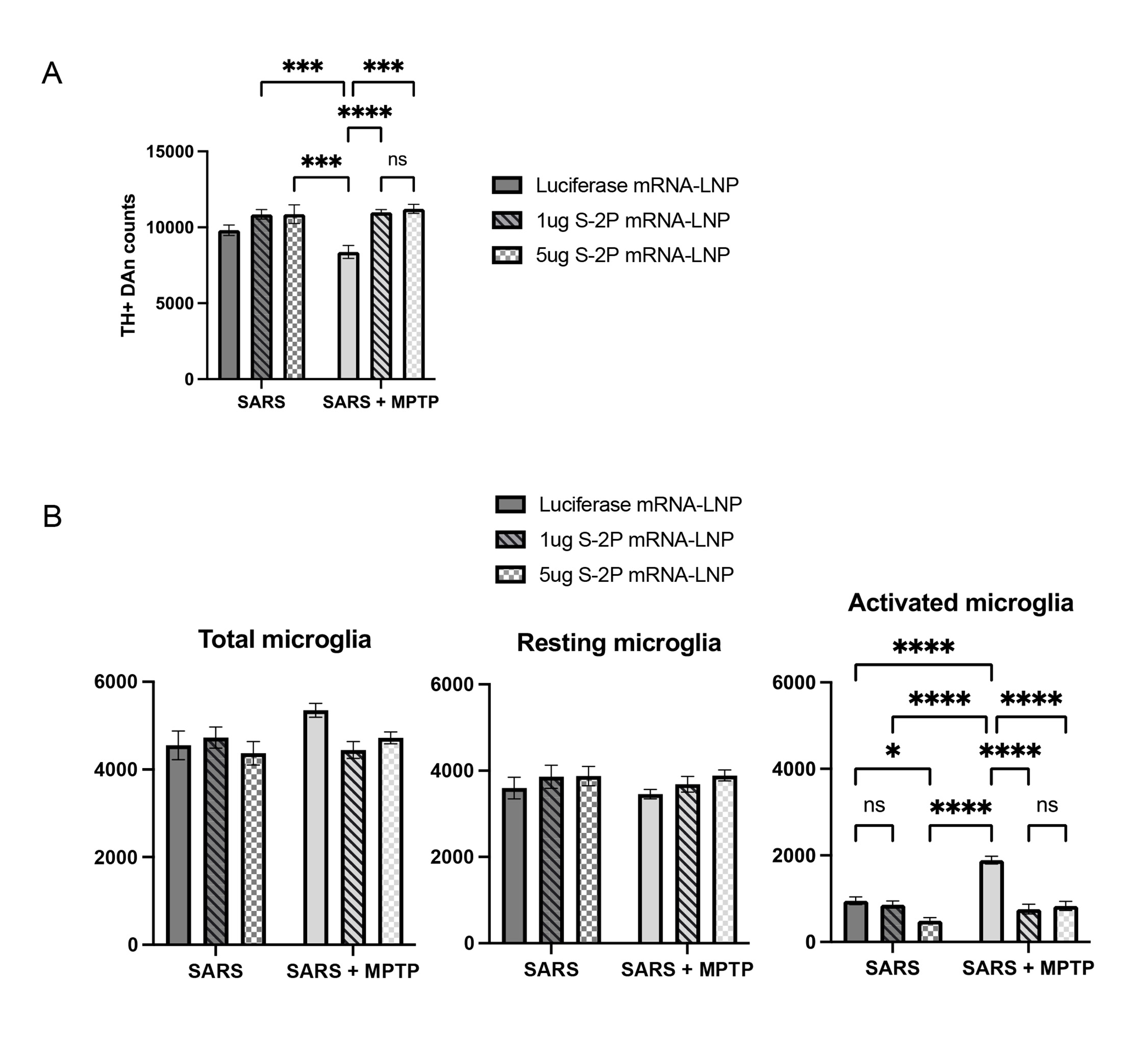

### Suppl fig 6

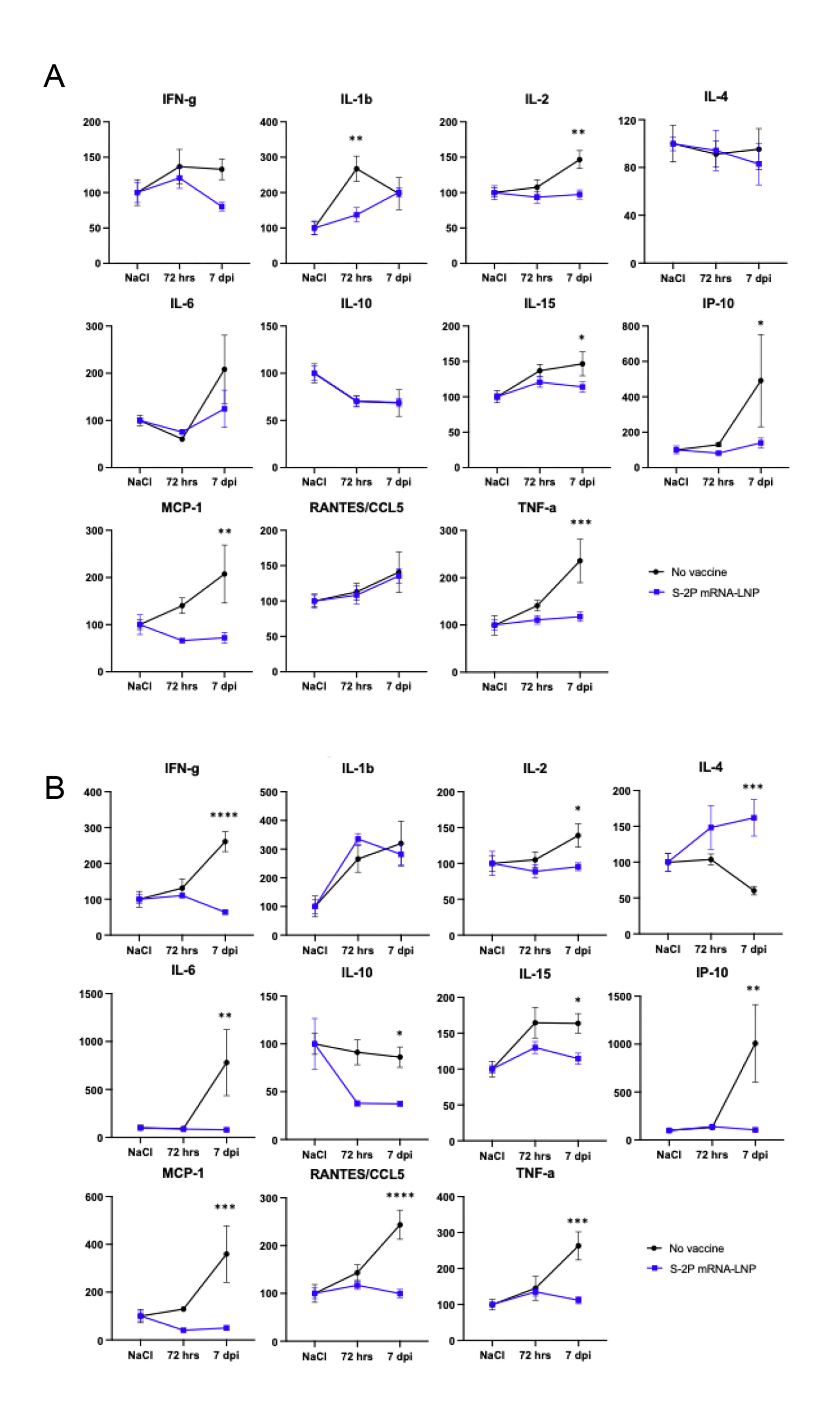
